# Customizing Multifunctional Neural Interfaces through Thermal Drawing Process

**DOI:** 10.1101/2021.05.17.444577

**Authors:** Marc-Joseph Antonini, Atharva Sahasrabudhe, Anthony Tabet, Miriam Schwalm, Dekel Rosenfeld, Indie Garwood, Jimin Park, Gabriel Loke, Tural Khudiyev, Mehmet Kanik, Nathan Corbin, Andres Canales, Alan Jasanoff, Yoel Fink, Polina Anikeeva

**Affiliations:** Research Laboratory of Electronics, Massachusetts Institute of Technology, Cambridge, MA, 02139, USA; McGovern Institute for Brain Research, Massachusetts Institute of Technology, Cambridge, MA, 02139, USA; Harvard/MIT Health Science & Technology Graduate Program, Cambridge, MA, 02139, USA; Department of Chemistry, Massachusetts Institute of Technology, Cambridge, MA, 02139, USA; Department of Chemical Engineering, Massachusetts Institute of Technology, Cambridge, MA, 02139, USA; Koch Institute for Integrative Cancer Research, Massachusetts Institute of Technology, Cambridge, MA, 02139, USA; Department of Biological Engineering, Massachusetts Institute of Technology, Cambridge, MA, 02139, USA; Department of Materials Science & Engineering, Massachusetts Institute of Technology, Cambridge, MA, 02139, USA; Kinetik Therapeutics LLC, Newton, MA, 02459, USA; Advanced Silicon Group, Lowell, MA, 01854, USA; Department of Brain & Cognitive Sciences, Massachusetts Institute of Technology, Cambridge, MA, 02139, USA; Department of Nuclear Science & Engineering, Massachusetts Institute of Technology, Cambridge, MA, 02139, USA; Advanced Functional Fabrics of America, Cambridge, MA, 02139 USA; Institute for Soldier Nanotechnologies, Massachusetts Institute of Technology, Cambridge, MA, 02139 USA

**Keywords:** thermal drawing, fibers, multifunctional neural probes, MRI, microdrives

## Abstract

Fiber drawing enables scalable fabrication of multifunctional flexible fibers that integrate electrical, optical and microfluidic modalities to record and modulate neural activity. Constraints on thermomechanical properties of materials, however, have prevented integrated drawing of metal electrodes with low-loss polymer waveguides for concurrent electrical recording and optical neuromodulation. Here we introduce two fabrication approaches: (1) an iterative thermal drawing with a soft, low melting temperature (T_m_) metal indium, and (2) a metal convergence drawing with traditionally non-drawable high T_m_ metal tungsten. Both approaches deliver multifunctional flexible neural interfaces with low-impedance metallic electrodes and low-loss waveguides, capable of recording optically-evoked and spontaneous neural activity in mice over several weeks. We couple these fibers with a light-weight mechanical microdrive (1g) that enables depth-specific interrogation of neural circuits in mice following chronic implantation. Finally, we demonstrate the compatibility of these fibers with magnetic resonance imaging (MRI) and apply them to visualize the delivery of chemical payloads through the integrated channels in real time. Together, these advances expand the domains of application of the fiber-based neural probes in neuroscience and neuroengineering.

## Introduction

Signaling complexity within the nervous system suggests the need for integration of bi-directional electrical, optical, and chemical interfaces within neural probes.^[1]^ Over the past decade, thermal drawing has enabled fabrication of versatile neural probes in the form of flexible miniature fibers that permitted multimodal neural interrogation while minimizing impact on local tissues.^[2–6]^ A typical fiber fabrication workflow begins with a centimeter-scale model, termed preform, which is produced by standard machining methods. The preform is then heated in a furnace above the glass transition temperature (T_g_) and melting temperature (T_m_) of its constituent materials and drawn into meters long microscale fibers.^[7]^ During this process, the preform dimensions are scaled down by several orders of magnitude, while preserving the overall cross-sectional geometry of the final fiber.^[8]^ This approach permits integration of multiple functionalities such as electrodes for recording of extracellular neuronal potentials, optical waveguides for optogenetic neuromodulation, and microfluidic channels for drug and gene delivery.

This one-step, top-down fabrication process can yield hundreds of probes from a single preform, and is a cost-efficient and scalable alternative to lithographic approaches that require resource-intensive cleanroom microfabrication techniques.^[9,10]^ Despite these advantages, widespread adoption of fiber-based probes by the neuroscience community has been hindered by several limitations.

Although thermal drawing is compatible with a wide range of metals, semiconductors, and polymers, recording electrodes previously incorporated into fibers were limited to either carbon-based polymer composites or drawable metals such as tin.^[2,11]^ Polymer composites exhibit limited conductivity due to the constraints on the weight fraction of conducting additives that can be incorporated into a polymer matrix without adversely modifying its T_g_ and viscosity.^[4,11]^ This poor conductivity restricts the electrode lateral dimensions (>∼20 μm) and fiber lengths (≲4 mm), which in turn limits the spatial resolution of extracellular electrophysiology ^[12]^ and increases the device footprint.^[4,11]^ Moreover, the heterogeneous volumetric distribution of filler particles in carbon composites introduces additional variability in the conductivity of polymer electrodes.^[4,11]^ Although metals can be thermally drawn, their T_m_ dictates the need for comparable T_g_ of the polymer cladding. For instance, while tin permitted integration of 5 μm electrodes into fiber-based probes, the necessary use of high T_g_ polymers such as poly(etherimide) and poly(phenylsulfone) precluded integration of low-loss optical waveguides due to the strong absorption of visible light in those materials.^[2]^ In addition to the materials limitations, the labor-intensive interfacing of each functional feature embedded within a fiber to the backend hardware has been the rate-limiting fabrication step and a barrier for broad adoption of fiber-based neural probes.

To overcome materials selection and interface challenges associated with fiber-based probes, we have implemented two approaches: (1) iterative thermal drawing with a soft, low T_m_ metal (indium); (2) convergence-drawing with a traditionally non-drawable metal (tungsten). These methods deliver multifunctional polymer fibers with low-impedance metal electrodes, low-loss polymer waveguides and microfluidic channels, while substantially reducing the complexity of back-end connectorization. We characterize the electrical, optical, fluidic and mechanical properties of these fibers and validate them *in vivo* for studies in mice and rats. Furthermore, we integrate multifunctional fibers into a light-weight and compact mechanical microdrive that allows fine electrode positioning post-implantation to maintain high-fidelity, single-neuron depth-specific recordings in freely moving mice. Finally, we demonstrate that these fibers elicit negligible artifacts in magnetic resonance imaging (MRI), and couple them with functional MRI (fMRI) to track the spatiotemporal kinetics of fluid delivery in the rat brain. This suggests that that fiber-based multifunctional probes could in future be leveraged for correlating global brain-wide activity to local neuromodulation.^[13–15]^

## Results and Discussion

### Fabrication of multifunctional neural probes through iterative and convergence thermal drawing

To enable scalable fabrication of neural probes with customizable functionalities & geometries, we developed two approaches: (1) Two-step iterative thermal drawing with soft, low T_m_ metal indium; and (2) convergence drawing with tungsten microwires. A two-step iterative thermal drawing approach is schematically illustrated in **Figure 1a**. The macroscale preforms of each modality (electrodes, optical waveguides, and microfluidic channels), termed here as the *first-step preforms*, are separately drawn into millimeter-scale *first-step fibers*. These first-step fibers are then arranged within the *second-step preform*. Drawing this second-step preform yields the microscale *second-step fiber*. This iterative size reduction approach using multi-step drawing enables the fabrication of microscale electrodes while preventing emergence of capillary instabilities.^[2,16,17]^

**Figure 1:**
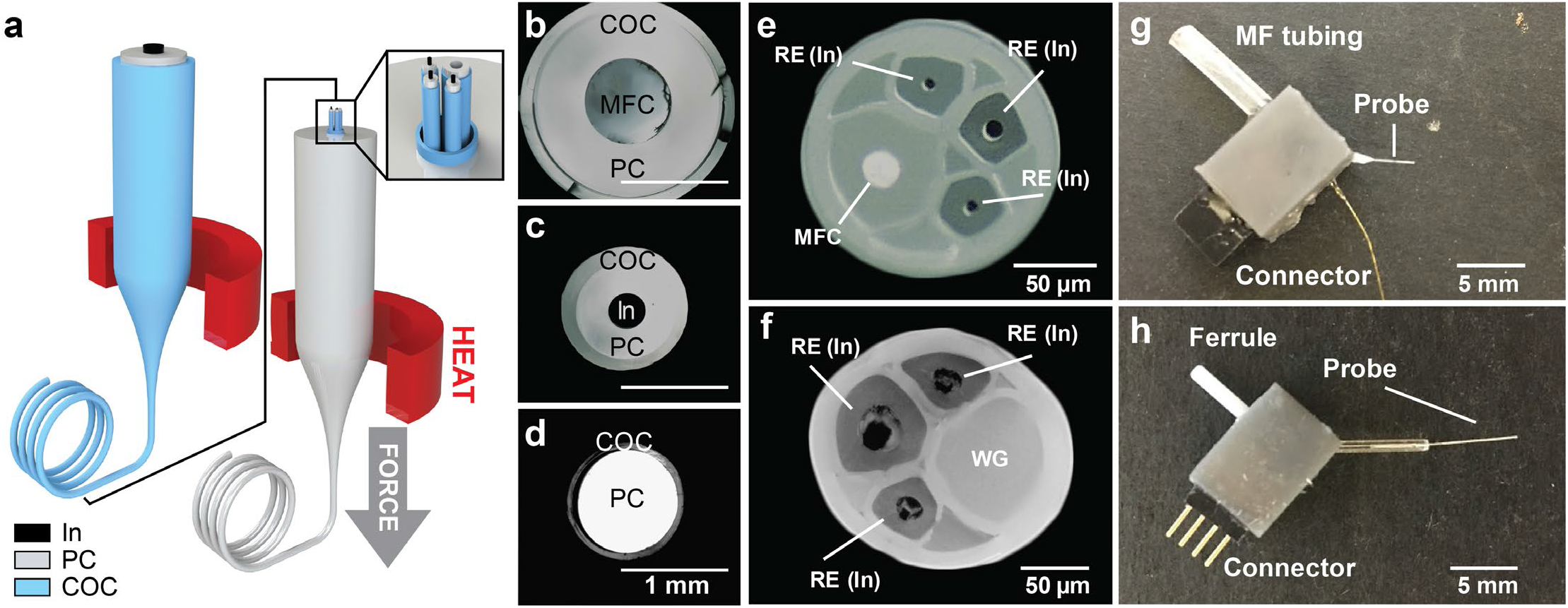
Multifunctional neural probe fabrication through the two-step thermal drawing process. (a) Schematic of the two-step TDP. (b-d) Cross-sectional microscope image of the fibers produced by the first iteration of the TDP. (e-f) Cross-sectional microscope image of the fibers produced by the second iteration of the TDP. (g-h) Final connectorized device. COC: cyclic olefin copolymer, MFC: microfluidic channel, PC: polycarbonate, In: indium, RE: Recording electrode.

Here, the first-step fibers consist of a recording electrode made of indium (In) encapsulated in polycarbonate (PC) and cyclic olefin copolymer (COC) cladding (**Figure 1b**), an optical waveguide with a PC core and COC cladding (**Figure 1c**), and a microfluidic channel made of a hollow PC/COC cladding (**Figure 1d**). Unlike other high T_m_ metals/alloys^[2,18]^, a low T_m_ of ∼156 °C enables indium to be co-drawn with an optically transparent PC/COC waveguide (PC refractive index n_PC_= 1.59, T_g_ = 150 °C; COC n_COC_ = 1.53, T_g_ = 158 °C).^[2,4,19]^ The macroscale first-step fibers are then arranged inside a COC/PC polymer cladding to form the second-step preform which is then drawn into a microscale fiber. As these first-step fibers can be arranged into any desired configuration, this “mix-and-match” approach enables straightforward customization of fibers for a given application. We illustrate this concept in two fiber designs: (i) an electro-fluidic fiber; and (ii) an opto-electric fiber. **Figure 1e** shows the cross-section of the electro-fluidic fiber (∼141.8 ± 7.4 μm) which comprises three indium electrodes (∼9.3 ± 2.1 μm) for electrophysiological recording and a hollow channel for concomitant delivery of fluids (19.1 ± 0.8 μm). Similarly, **Figure 1f** depicts the cross-section of the opto-electric fiber (∼162.2 ± 6.1 μm) that enables simultaneous light delivery and electrical recording, and features three indium electrodes (∼21.6 ± 7.8 μm) and a PC/COC optical waveguide (∼67.3 ± 5.5 μm). **Figure S1** demonstrates another example of the iterative drawing method to achieve a more complex fiber geometry. Here, multifunctional first-step fibers are fabricated (**Figure S1a**,**b**), and stacked into a linear pattern before being drawn again into a microscale fiber (**Figure S1c**,**d**). By cutting the second-step fiber at an angle (**Figure S1e**), or in a step-wise manner (**Figure S1f**,**g**), depth specific recording and modulation can potentially be achieved with this design.

The iterative fiber drawing approach not only enables customized fiber design, but also facilitates the back-end connectorization process, which transforms a fiber segment into a fully functional neural probe **(Figure S2)**. Drawing of the second step preform without prior consolidation allows for facile separation of individual components in a second step fiber upon removal of its outer COC cladding due to strain **(Figure S2)**. The mechanical etching of the outermost sacrificial cladding exposes individual functional components (electrodes, waveguides, channels) in the fiber and provides independent access to each component. The electrode microfibers are then connected to header-pins using a conductive adhesive, the waveguide is coupled to a ceramic ferrule, and the hollow channel is connected to the plastic tubing for fluid delivery. The sacrificial PC cladding from the remaining exposed fiber (implantable front end) is chemically etched. Encapsulation of the backend input/output (I/O) hardware inside a 3D printed microdrive shuttle completes the overall fabrication process. **Figure 1g** and **1h** show images of fully connected fiber-based devices. Through this approach, the connectorization process has been reduced to 9 steps (∼1h per device), compared to 18 steps (∼4h per device) used in previous studies.^[4]^

Another approach to overcome materials constraints in multifunctional fiber-probe design relies on recently demonstrated convergence drawing.^[20]^ This approach overcomes constraints on the preform components to have similar glass transition temperatures (T_g_) and melt viscosities at the drawing temperature.^[6,8]^ During convergence drawing, a microwire of a material with T_m_ (or T_g_) significantly higher than the drawing temperature is fed into a hollow channel within the preform, which collapses and converges the wire into the polymer cladding of the resulting fiber (**Figure 2a**). This approach enables incorporation of high conductivity metallic electrodes independent of their T_m_ and thereby widens the palette of functional materials that can be integrated into fibers.^[6,8]^ To illustrate the versatility of this fiber drawing method we fabricate two different fiber designs: (i) a tetrode-like fiber; (ii) a multifunctional fiber with electrical, optical, and microfluidic features. First, we fabricate a 108.94 ± 9.4 μm PC (T_g_ = 150 °C) fiber comprising four tungsten electrodes arranged in a tetrode like configuration (**Figure 2b**). The preform is produced by machining four grooves (1 mm) into a PC rod (1/8 inch in diameter), which is then wrapped with PC sheets and consolidated in a vacuum oven (Methods) to obtain a final diameter of ∼4 mm. The preform is then drawn into ∼50 m long fiber while simultaneously feeding tungsten microwires into the hollow channels. The resulting microscale fiber has four closely packed 25 μm electrodes that are separated by 10-12 μm (**Figure 2b**). This process can be easily extended to a higher number of electrodes, either through a convergence of N electrodes (N>4) or a subsequent convergence of multiple tetrode fibers (N × 4-electrodes) akin to iterative thermal drawing described above.

**Figure 2:**
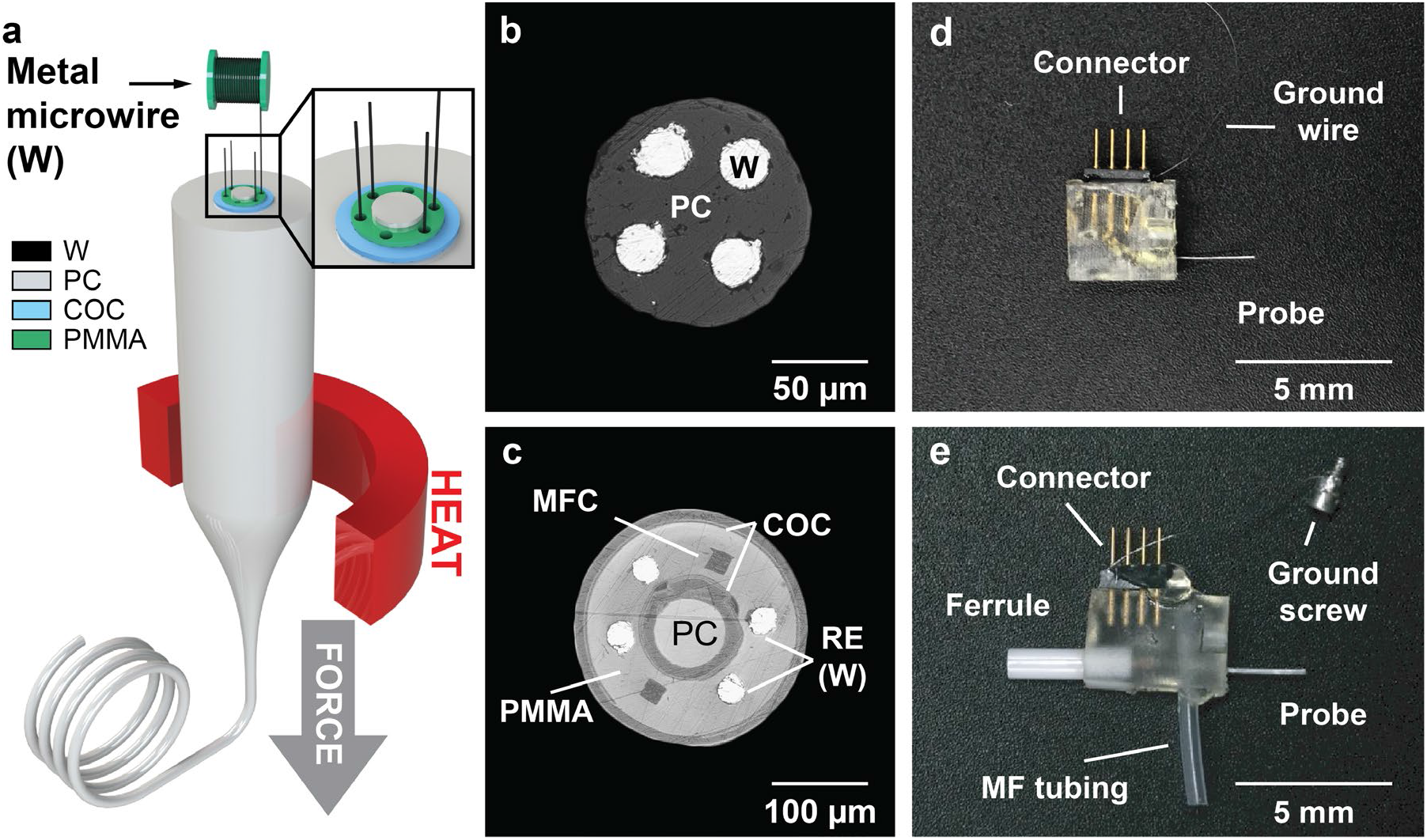
Multifunctional neural probe fabrication through the convergence thermal drawing process. (a) Schematic of the convergence drawing process. (b-c) Cross-sectional microscope image of the fibers produced by the convergence TDP. (d-e) Final connectorized device. COC: cyclic olefin copolymer, PMMA: Polymethyl methacrylate, PC: polycarbonate, W: tungsten, MFC: microfluidic channel, RE: Recording electrode.

Following demonstration of integrating tungsten electrodes into a transparent polymer PC, we apply convergence drawing to create a multifunctional probe that features four tungsten recording electrodes, a PC/COC optical waveguide, and two microfluidic channels (**Figure 2c**). The preform for this fiber was produced by standard machining, similar to fibers described above, and then drawn into ∼100 m long fiber with a 40-45-fold reduction in its cross-sectional dimensions. Four 25µm tungsten microwires were simultaneously fed into the fiber during drawing (**Video S1**). The overall cross-sectional diameter of the fiber was 225.50 ± 8.7 μm (∼350 μm prior to etching of the sacrificial layers). The final linear dimensions of the waveguide core and microfluidic channels were 108 ± 4.9 µm and 19.27 ± 1.5µm, respectively. (**Figure 2c)**. Similar to the iterative drawing, convergence drawing simplifies the interfacing to the back-end connectors. **Figure S3** schematically depicts a typical process flow, wherein solvent etching exposes metal microwires and/or the PC/COC waveguide, which are subsequently connected to electrical pins and optical ferrules, respectively. The microfluidic channels are accessed through an orthogonal T-connection and the entire assembly is encapsulated into a 3D printed shuttle drive. Complete, ready for implantation tetrode-like and multifunctional fibers (**Figure 2c,d**) can be produced in 4-6 steps (30 min). Thus, the iterative drawing and convergence drawing approaches presented in this work not only expand the array of materials used in fiber-based probes, but also address the bottleneck associated with back-end interfacing of these devices.

### Characterization of optical, electrical, mechanical, and fluidic properties of fiber-based probes

We next characterized electrical, optical, fluidic, and mechanical properties that will dictate the performance of the fiber-based probes *in vivo*. Electrochemical impedance spectroscopy was employed to assess the utility of the integrated tungsten and indium microelectrodes for neural recording (**Figure 3a**). The average electrode impedance was found to be in the range of 71.8 ± 29.8 kΩ for tungsten and 338.1 ± 167.8 kΩ for indium at 1 kHz, which is comparable to state-of-the-art recording probes featuring metallic electrodes with similar dimensions.^[1,12]^ Moreover, the inter-electrode impedance through the polymer cladding was found to be >2 MΩ at 1 kHz which indicates negligible inter-electrode cross talk (**Figure S4**).^[21]^ Tungsten electrodes are routinely used for *in vivo* electrophysiology and are known to form stable interfaces with the surrounding tissue.^[22,23]^ However, little is known about the nature of the electrochemical interface between indium and the neural tissue. From the established Pourbaix diagram for indium (**Figure S5a**),^[24]^ we hypothesized the presence of a self-passivating native oxide layer on the surface of indium microelectrodes at physiological conditions (pH 7.4, E= 0V). This was indeed corroborated by the presence of a reversible In^2+^/In^3+^ redox peak in the –0.9 to –0.8V potential window (vs. Ag/AgCl) which is observed for both macro- and microscale indium electrodes (**Figure 3b**). This self-passivating native oxide layer on the electrode surface can be beneficial to establishing a capacitively dominant electrochemical interface that minimizes faradaic charge transfer reactions.^[25]^ To further quantify this observation, the Nyquist plot for indium was fitted to a Randle’s equivalent circuit and the corresponding charge transfer resistance (R_ct_) value was calculated (**Figure 3c**). A high value of R_ct_ (4.68 MΩ) suggests slow charge-transfer reactions at the metal/electrolyte interface, and thus a predominantly capacitive interface for indium electrodes.

**Figure 3:**
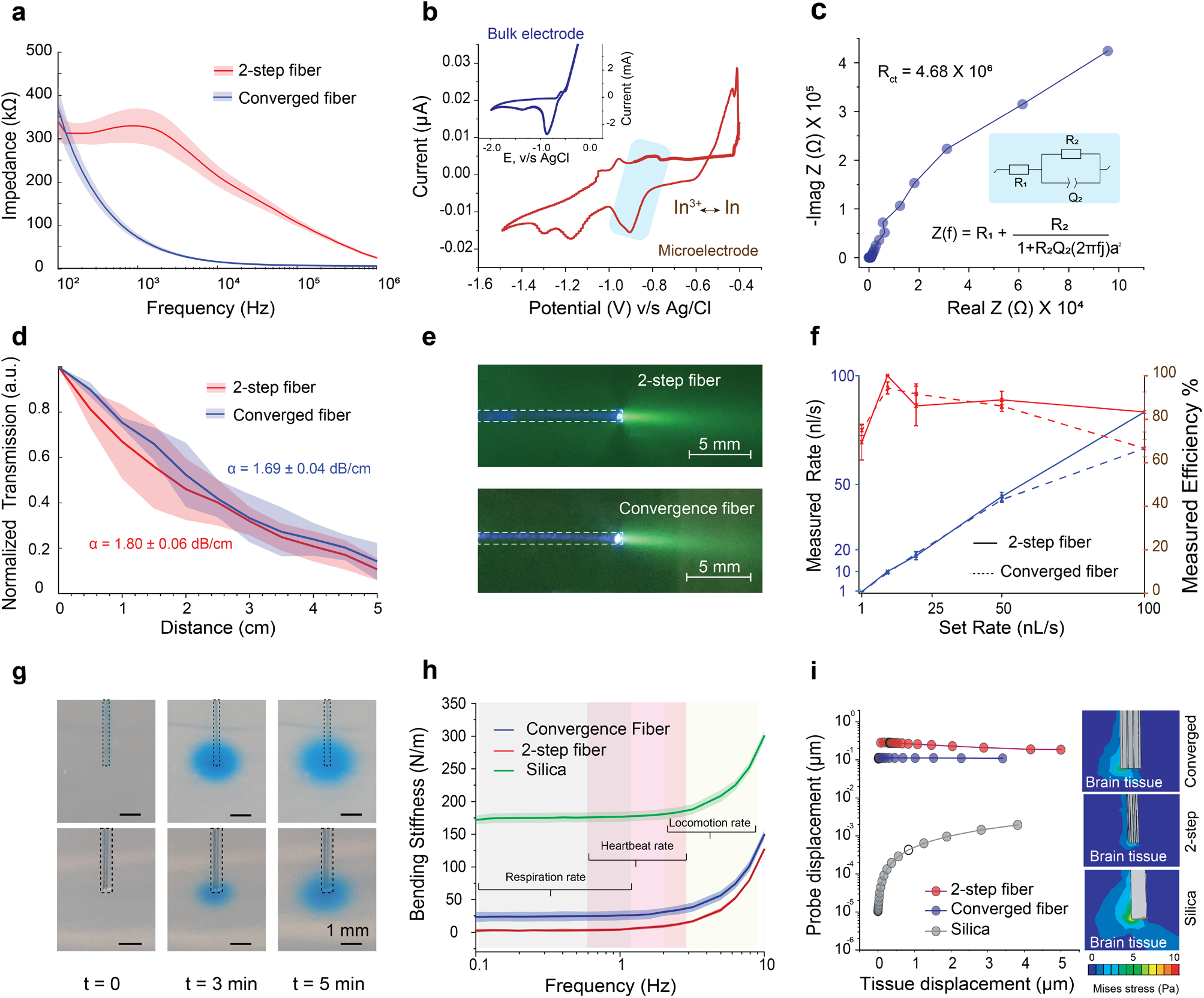
Characterization of the two-step and converged multifunctional fibers. (a) Electrochemical impedance spectroscopy of the two-step and converged fiber probes. (b) Cyclic voltammogram in 1xPBS of bulk indium and indium microelectrode from neural probe of fig. 1e. (c) Nyquist plot and equivalent circuit model for indium electrode in PBS (d) Optical loss characterization of the two-step and converged fiber probes. Shaded areas represent s.d (n=3). (e) Illustration of light emission profiles of waveguides via coupling of 473nm laser in a diluted fluorescein solution. (f) Output injection rate and injection efficiency of the microfluidic channel of two-step and converged fiber probes. (n = 4, 2 cm long). (g) Illustration of microfluidic capabilities via infusion of a blue dye at 100nl/min into a phantom brain (0.6% agarose gel). (h) Bending stiffness measured for two-step fiber, convergence fiber, and a 300 µm silica optical fiber. Shaded areas represent s.d (n=3). (i) The displacement for two-step fibers, converged fibers and 300μm silica fibers during 0-5 μm lateral displacement of the brain tissue (left); Mises stress profile within the brain tissue for two-step fibers, converged fibers and 300 μm silica fibers at 5 μm lateral deformation of the brain tissue.

We then characterized the integrated waveguides within the two-step and converged fibers. A transmission loss coefficient (α) of 1.8 ± 0.06 and 1.69 ± 0.04 dB/cm at 473 nm was determined for the opto-electric and the multifunctional fibers, respectively (**Figure 3d**). The obtained values are comparable to those of previously reported multifunctional fibers, and indicate that these integrated polymer waveguides are suitable for optogenetic modulation mediated by blue-light activated opsins *in vivo*.^[2,4,26]^ **Figure 3e** shows typical illumination profiles from the waveguides for fibers produced via iterative and convergence drawing methods immersed in a solution of Fluorescein dye (0.01 mM).

Next, we quantified the efficiency of fluid delivery by infusing Evans’ Blue dye solution at the injection rates that are commonly used for intracerebral infusions (**Figure 3f**). An efficiency between 65 –100% was observed for injection rates between 1-100 nL/s for both the electro-fluidic and multifunctional fibers. **Figure 3g** shows a typical time-course of dye injection in a phantom brain (0.6% agarose gel).

Finally, we characterized the mechanical properties of both fibers by quantifying their bending stiffness in a single cantilever mode (**Figure 3h**). We found that fibers produced by two-step and convergence drawing showed substantially lower bending stiffness than a conventional silica fiber of comparable dimensions (300 µm) across the entire frequency spectrum corresponding to respiration, heartbeat and locomotion rates (0.1–10 Hz). A decrease in bending stiffness positively correlates with reduced relative micromotion between brain tissue and implanted probes, which is anticipated to reduce the tissue damage and facilitate their chronic use.^[27]^ The two-step fibers were found to have 10-12 times lower stiffness values compared to converged multifunctional fibers due to their smaller dimensions (140 μm vs 225 μm) and substantially lower Young’s modulus for indium (12.74 GPa) as compared to tungsten (340 GPa).^[28]^ The stress distribution in the polymer cladding visualized via finite elemental analysis (FEA) for a single cantilever bending mode qualitatively corroborates the experimentally observed trends of stiffness values (**Figure S5b**).

We next modeled the relative micromotion between the implanted fiber probes and the brain tissue using FEA. From the displacement plots, it is apparent that relative micromotion between the implant and the tissue is higher for mechanically rigid silica fibers as compared to compliant polymer fibers (**Fig. 3i (left))**. The deflection at the implant tip for silica fibers is predicted to be ∼2 – 4 orders of magnitude lower than that for the flexible polymer fibers. The FEA also captures the difference in micromotion between the two-step and converged fibers, wherein the soft indium-containing fibers experience larger displacement (and hence smaller relative motion) than the stiffer converged fibers. The distribution of von-Mises stress profiles around the implant tips produced by this relative micromotion is in agreement with the bending-stiffness trends experimentally observed for the respective probes (**Fig. 3i, right)**. Note, that depending on the application, the bending stiffness can be further adjusted by reducing the fiber dimensions and selecting softer polymers as cladding.

### Recording and stimulation of neural activity in vivo using two-step and converged fiber probes

To evaluate our probes’ ability to chronically record spontaneous neuronal activity, we implanted the two-step electro-fluidic fibers (**Figure 1e**) and the converged multifunctional fibers (**Figure 2c**) into the medial prefrontal cortex (mPFC) of wild type (WT) C57BL/6 mice. The central role that mPFC plays in cognitive and executive processes has made this brain region a target of numerous neuromodulation studies, offering a rich field for comparison of our probes’ performance to prior observations.^[29–31]^

Both designs reliably recorded extracellular potentials for at least four weeks following implantation surgeries, and single unit activity was routinely detected over this time period. Similar to prior studies with Sn microelectrodes,^[2]^ electrophysiological recording with indium electrodes in the two-step fibers had a signal to noise ratio (SNR) of 7.4 ± 0.7 and a noise floor of 114.4 ± 39.3 µV peak-to-peak at one week (**Figure 4a**), and a SNR of 10.9 ± 2.8 and noise floor of 94.5 ± 21.1 µV at four weeks post-implantation (**Figure 4e**). Principal component analysis (PCA) followed by gaussian mixture model (GMM) clustering ^[32]^ isolated 2 putative single units at week 1 (**Figure 4b**,**c**). For these units, cluster quality was confirmed as intermediate by calculating L-ratio (<0.1) and isolation distance (>50).^[33]^ Although one of the isolated units was lost during subsequent recordings, the second unit was preserved up to week 4 (**Figure 4e**,**f**,**g**) and likely up to week 8 (**Figure S6**) as indicated by its stable average waveform (**Figure 4c**,**g**) and inter-spike interval (ISI) (**Figure 4d**,**h**). ^[34]^

**Figure 4:**
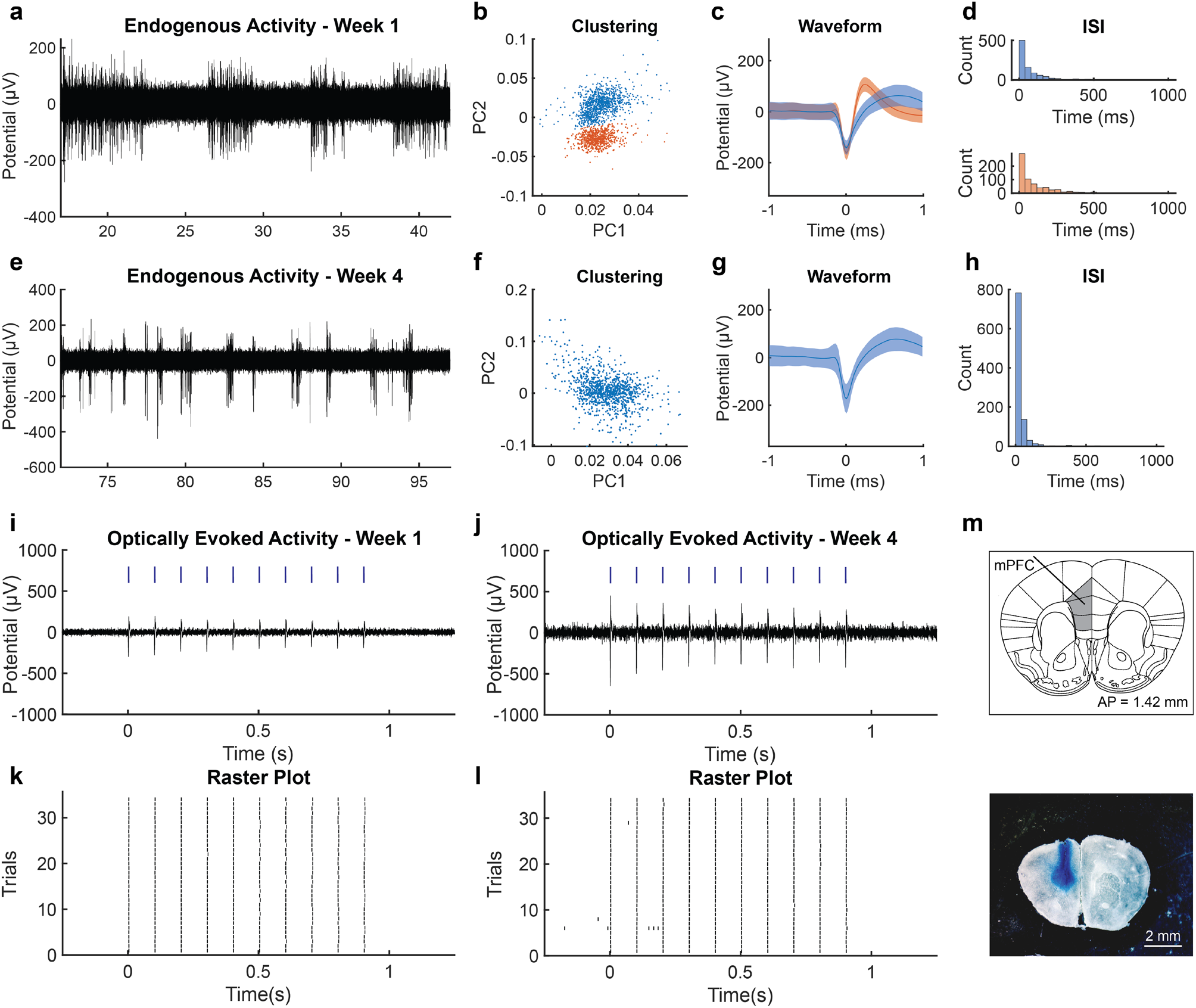
Two-step opto-electric and electro-fluidic neural probes enables chronic recording of endogenous and optically-evoked neural activity *in vivo*. (a) (a-h) Endogenous activity recorded using the two-step fiber. (a) endogenous activity 1 week after implantation containing 2 separable single-unit activity. (SNR = 6.88, noise floor = 89 µV) (b) Principal-component analysis (PCA) of the putative unit detected in (a). (c) Average waveform (STA) shows the waveform of the separated units. (d) Interspike interval of each separated unit. (e-h) similar to (a-d) endogenous activity, PCA, waveform and ISI for week 4 after implantation. (SNR = 12.91, noise floor = 80 µV) A single unit detected and is similar to the one of the unit from week one. (i-l) Simultaneous optogenetic stimulation (wavelength λ=473 nm, 5 ms pulse width, 10 Hz, 1s-long pulse train, power density 10 mW/mm^2^, blue markers indicate laser pulse. Activity recorded 1 week (i) and 4 weeks (j) post implantation. raster plot of detected evoked potentials over 31 stimulation trials (k,l) highlight the reproducibility of the optically-evoked activity. (m) micrograph of brain slices following Evans’ Blue dye injection (2% in NaCl, 2µl over 20mn) through the microfluidic channels and transcardial perfusion of the animal.

Similarly, tungsten electrodes in the converged multifunctional fibers also enabled weeks-long tracking of single units. These recordings yielded a SNR values of 15.6 ± 5.2 and 12.4 ± 2.4 and a noise floor 36.2 ± 8.4 µV and 41.5 ± 2.4 µV at weeks 2 and 3, respectively (**Figure 5c,g**). Two single units were isolated and tracked from week 2 (**Figure 5a,b,c,d**) to week 3 (**Figure 5e,f,g,h**) (L-ratio <10^−9^, isolation distance >105), as indicated by the stable waveform and ISI.

**Figure 5:**
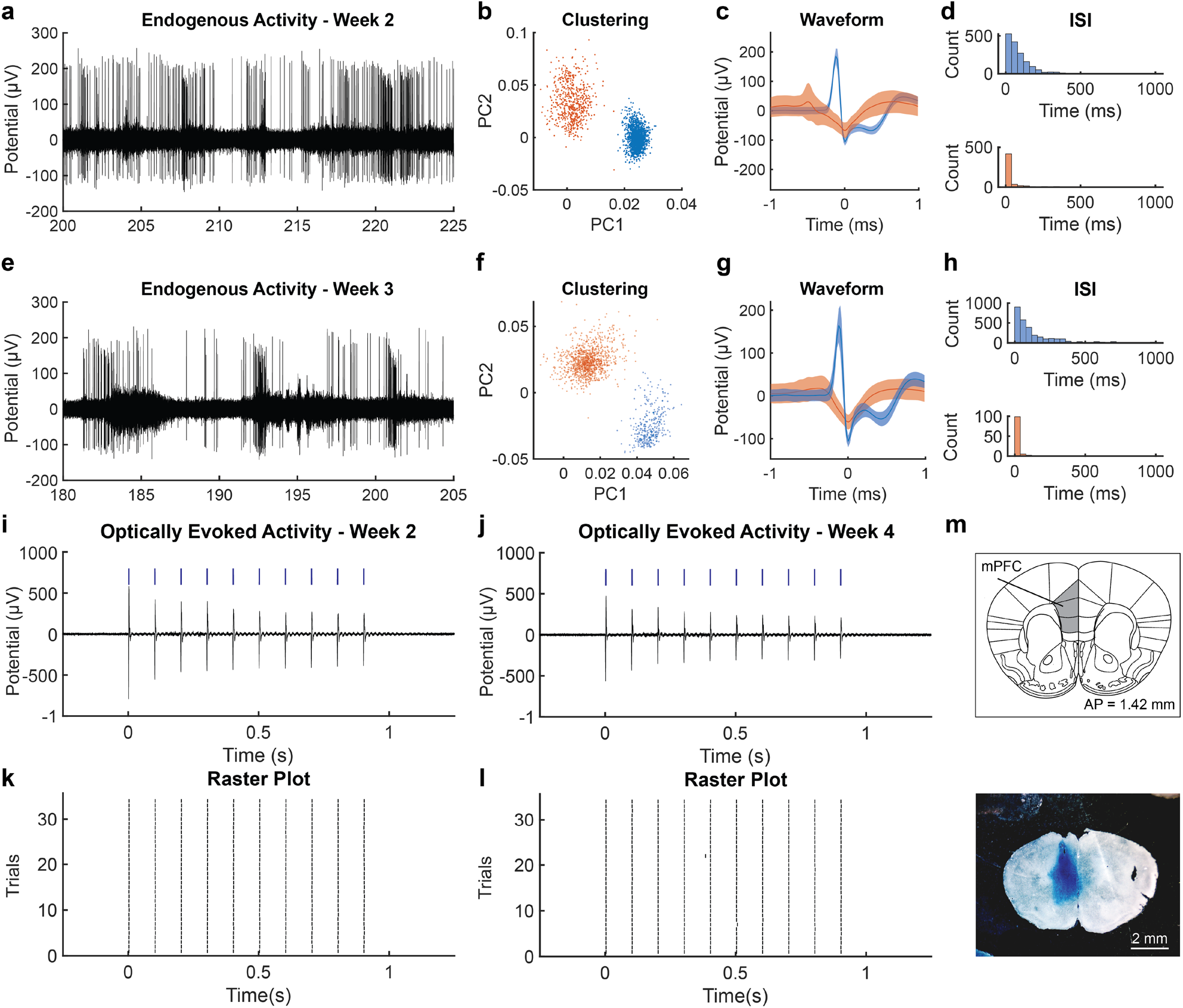
Converged multifunctional neural probes enables chronic recording of endogenous and optically-evoked neural activity *in vivo*. (a) (a-h) Endogenous activity recorded using the convergence fiber. (a) endogenous activity 2 week after implantation containing one separable single-unit activity and one multi-unit activity. (SNR = 21.26, noise floor = 46 µV) (b) Principal-component analysis (PCA) of the putative unit detected in (a). (c) Average waveform of the separated units. (d) Interspike interval of each separated units. (e-h) similar to (a-d) endogenous activity, PCA, waveform and ISI for 3 weeks after implantation (SNR = 14.07, noise floor = 40 µV). Both multi-unit and single unit detected appear similar to week 2. (i-j) Simultaneous optogenetic stimulation (wavelength λ=473 nm, 5 ms pulse width, 10 Hz, 1s-long pulse train, power density 10 mW/mm^2^, blue markers indicate laser pulse. Activity recorded 2 weeks (i) and 4 weeks (j) post implantation. Raster plot of detected evoked potentials over 31 stimulation trials (k,l) highlight the reproducibility of the optically-evoked activity. (m) micrograph of brain slices following Evans’ Blue dye injection (2% in NaCl, 2 µl over 20mn) through the microfluidic channels and transcardial perfusion of the animal.

Next, we tested the ability of the two-step and converged probes to record neural activity during optical stimulation. To sensitize the neurons to light, we first injected a viral vector carrying a transgene that encodes for a blue light-sensitive cation channel channelrhodopsin-2 (ChR2) under the excitatory neuronal promoter calmodulin kinase II α-subunit (*AAV5-CaMKIIα::ChR2-EYFP)*^[35]^ into the mPFC of WT mice. Following a 4-weeks incubation period, we implanted either the opto-electronic two-step fiber (**Figure 1f**) or the converged multifunctional fiber (**Figure 2c**) at the injection coordinates (Methods). For both designs, we reliably recorded electrophysiological activity evoked by optogenetic stimulation (λ = 473 nm, 10 Hz, 10 mW/mm^2^, 5 ms pulse width) (**Figure 4i,j** and **Figure 5i,j**). The recorded activity was correlated with the onset of the laser stimulation (Jitter = 0.6 ms, peak latency = 3.7 ms), and reproducibility of the response was qualitatively assessed with the spike raster plot (32 stimulation trials, 5-s inter trial interval) (**Figure 4k,l** and **Figure 5k,l**). To further confirm the physiological origin of the optically-evoked potentials and to rule out photo-electrochemical artifacts,^[36]^ we performed an array of control experiments. We observed that violet light (λ = 420 nm, 4 mW/mm^2^, 10 Hz, 5 ms pulse width) pulses (**Figure S7a,c**) and but not green light (λ = 565 nm, 4 mW/mm^2^, 10 Hz, 5 ms pulse width) pulses (**Figure S7b,d**) elicited an electrophysiological response. Additionally, contrary to mice transduced to express ChR2 (**Figure S7e,g**), mice transduced with *AAV-CamKIIα::EYFP* (a control virus carrying only a gene for an enhanced yellow fluorescent protein) did not exhibit any optically-evoked activity (**Figure S7f,h**). We also observed that 100 Hz stimulation pulse train (**Figure S7j,l**) evoked neuronal activity that was uncorrelated with the optical pulses, which is consistent with the ChR2 kinetics.^[37]^

The neuronal activity was further correlated to a previously observed behavioral response to optical stimulation in the M2 area of the mPFC.^[4,38]^ The mice were subjected to an open field test divided into three epochs OFF/ON/OFF, each lasting 3 mins. During the ON epochs, mice received optical stimulation (473nm, 20Hz, 10ms, 30mW/mm^2^) and no stimulation was applied during OFF epochs (**Figure S8a,b**). Mice expressing ChR2 exhibited a higher locomotor activity during stimulation epochs as compared to pre- and post-stimulation epochs, while the control group showed no effect of optical stimulation (**Figure S8c)**.

Finally, the ability of our probes to deliver chemical cargo *in vivo* through the integrated microfluidic channels was evaluated by delivering a bolus of Evans’s blue dye (2% in sterile NaCl solution, 2 µL, 20 nL/s) into the mPFC of WT mice. Delivery of the dye was then confirmed *via* optical microscopy of the extracted brain slices (**Figure 4m** and **Figure 5m**).

Together, these findings illustrate that the two-step and convergence techniques extend the utility of thermal drawing to additional classes of materials and permit fabrication of neural probes that record and stimulate neuronal activity across multiple modalities. While indium has not been commonly used in neural recording devices, our findings demonstrate its utility for recording endogenous as well as optically-evoked activity *in vivo* for at least eight weeks. Note that, indium electrodes exhibited higher impedances and lower SNRs compared to their tungsten counterparts integrated via convergence into polymer fibers. However, surface coating strategies such as electrochemical deposition of conducting polymers or noble metals could potentially further optimize their performance.^[39]^ Convergence drawing can also be extended beyond tungsten to microwires composed of platinum-iridium that can potentially enable additional features including electrical stimulation via multifunctional fibers.^[40]^

### Coupling fibers to microdrives: Fabrication and characterization

To enable depth-resolved interrogation of neural circuits, we adapted the custom-screw and shuttle drive mechanism developed by Voigts and colleagues ^[41]^ to a microdrive compatible with our flexible polymer fibers. Similar to the prior design, our microdrive is composed of four functional elements: the drive body, the shuttle that carries the neural probe, a ceramic retaining collar and a custom screw (**Figure 6a**). Vertical translation of the shuttle is actuated by rotating the screw (**Figure S9a-b**). Step-by-step assembly of the microdrive is illustrated in **Figure S10**.

**Figure 6.**
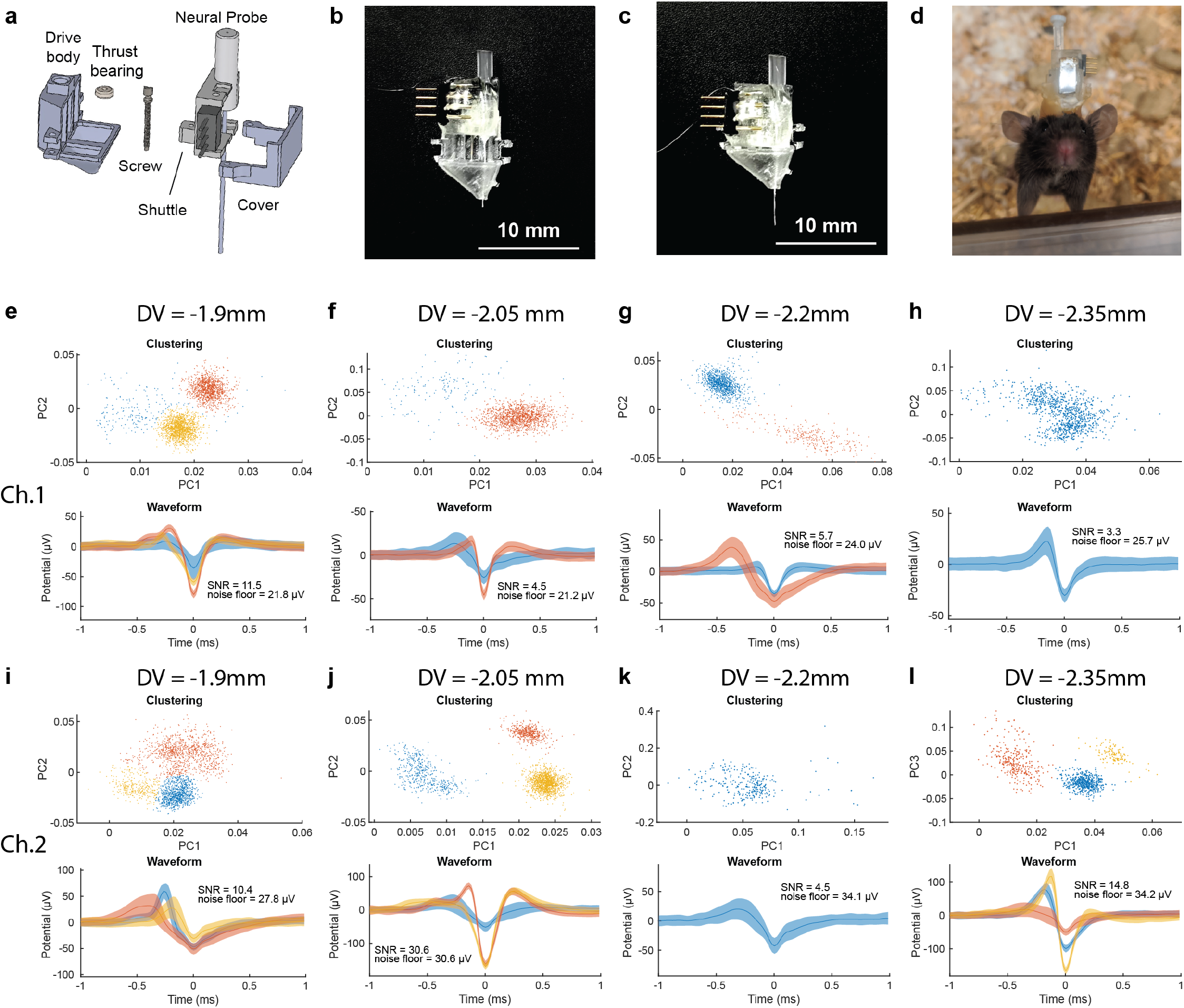
- Microdrive assembly enables depth-specific recording of endogenous neural activity *in vivo*. (a) Exploded view of the microdrive assembly composed of a drive body, a dental cement thrust bearing, a custom-made screw, a neural probe mounted on a shuttle, and a cover. (b-c) neural probe assembly onto the microdrive on the up (b) and down (c) position. (d) Picture of the assembly implanted into the mPFC of a C57BL/6 wild type mice. The implant was placed at AP= +1.7 mm, ML= 0.4 mm, DV= −1.9 mm during the first day, and was lowered by 150 µm (1 turn) every day for 3 days. Endogenous neural activity was successfully recorded at each depth for channel 1 (e-h) and 2 (i-l) and putative single unit activity were isolated by principal-component analysis (top), and the average trace illustrates the waveform of the separated units (bottom).

The resulting drive enables controlled linear motion with a travel range of 4 mm (**Figure S9c,d,e**) sufficient to probe the majority of the dorsoventral span in multiple regions in a mouse brain.^[42]^ The assembled microdrive has an overall maximum volume of 25 × 15 × 8 mm^3^ (excluding the implanted fiber), and a weight of 1g ± 0.02g (**Figure 6b,c,d**). Using an open-field test of the locomotor activity, no differences in speed and track length were identified between microdrive-carrying mice and naïve counterparts. As expected tethering the drives to external hardware produces modest yet significant decrease in locomotion motivating future implementation of wireless interfaces (**Figure S11**). To characterize the linear displacement of the fiber probes via the microdrives, we lowered them into a phantom brain (0.6% agarose) while tracking the position of the fiber tip under a microscope. The custom screw which has a thread pitch of 150 μm is expected to produce ∼150 μm fiber displacement for each counterclockwise turn.^[41]^ We found that the observed displacement of the fiber tip was consistently higher than the targeted value (**Figure S9d**). This tracking error could be attributed to the imprecision of the manual rotation of the screw. Conversely, the observed lateral deflection was found to be less than 30 μm per millimeter of vertical displacement, ensuring precise targeting in the brain (**Figure S9e**). We additionally validated the depth-specific microfluidic injection of the microdrive-fiber assembly by delivering a bolus of Evans’ Blue dye (2% into water, 500 nL, 100 nL/min) at 3 different drive positions (**Figure S9f**).

### Coupling fibers to the microdrive for depth specific in vivo electrophysiology

To test the utility of the microdrive-coupled multifunctional fibers to probe neural activity along the dorsoventral (DV) axis, we implanted the tetrode-like converged fiber (**Figure 2b**) into the mPFC of wild type mice. Fibers including tungsten electrodes were chosen over those employing indium due to the lower impedance and higher SNR associated with these probes. During implantation, the assembly was positioned slightly above the mPFC (AP +1.7 mm, DV −1.9 mm, ML ± 0.4 mm). Following a 3-day recovery period, we recorded endogenous electrophysiological activity under isoflurane anesthesia every day for 4 days. After each recording session, the microdrive screw was turned counterclockwise (1 turn) to advance the implant by 150 μm. The tissue was allowed to recover for a full day prior to the next recording session. Given that 150 μm distance exceeds the typical single-unit recording range of an electrode,^[43]^ each recording session was anticipated to yield a different set of units. We recorded spiking activity with SNR=10.7 ± 9.0 and noise floor = 27.4 ± 5.2 µV each day on two of the four channels **(#1 - Figure 6e-h, #2 - Figure 6i-l)**, while the other two channels recorded little to no spontaneous activity and were used as reference. We found that potentials could be isolated, and clustered into putative single units and multi-unit activity **(Figure 6e-l)**. For each isolated cluster, we measured its average firing rate, inter-spike interval (ISI), L-ratio, and isolation distance ^[33,34,44]^ (**Figure S12**). The differences in these parameters observed for each unit indicate that electrophysiological activity was recorded from distinct neurons upon daily advancement of the probes, while the L-ratio and isolation distance illustrate an intermediate to good quality of clustering for 10 out of 16 clustered units (L-ratio <0.01, isolation distance >20). Similar results were obtained for electrophysiological recording sessions in freely moving mice (**Figure S13**). While on average the SNR was lower and noise floor was higher than the values observed for recordings in anesthetized subjects (SNR = 8.3 ± 6.1, noise floor = 28.8 ± 5.4 µV), the majority of putative single-unit and multi-unit activity were recovered (15/18 units), with an additional unit being detected in the awake but not in the anesthetized animals (**Figure S13h**).

This study highlights how thermally-drawn neural probes can be integrated with compact and light-weight microdrives to enable recordings from several different loci along the DV axis, in both anesthetized and freely moving mice. Given that a similar design can be applied to any multifunctional fiber-based probe, this approach can be further exploited to perform multimodal depth specific interrogation of neural circuits using optical and chemical functionalities.

### In vivo MRI imaging in the presence of the multifunctional fibers

Pairing neural probes with magnetic resonance imaging (MRI) enables simultaneous local interrogation of neural ensembles and anatomical and functional characterization across the entire brain. This approach has been leveraged to understand neurophysiological correlates of functional MRI (fMRI) signals,^[13]^ brain wide circuit activation patterns in response to deep brain stimulation,^[15]^ or to verify an implant placement.^[45]^ However, the mismatch in the magnetic susceptibility between the typical neural recording electrodes and brain tissue manifests in large image artifacts making it challenging to visualize the surrounding brain structures. Here we evaluated the compatibility of both two-step and converged multifunctional fibers with MRI in a 9.4T small animal scanner. We applied fiber-based probes implanted in subcortical brain structures of rats to focally deliver a Gd^3+^-based MRI contrast agent ProHance (10mM) through the integrated microfluidic channels. During an imaging session, the implanted rat was placed in the MR scanner and the microfluidic channel was connected to an external infusion pump (**Figure 7a,b**). We observed that the converged fibers produced a minimal tissue distortion artifact while the two-step fiber exhibited an almost artifact-free profile (**Figure S14**). Artifact sizes of ∼400 μm and ∼200 μm were measured for converged and two-step fibers, respectively, which is comparable to their actual diameters of (225 ± 8.7 µm and 141.8 ± 7.4 µm respectively), indicating negligible loss of functional response visualization during MRI. We note that the artifacts created by polymer fibers are substantially smaller than those produced by standard monopolar stimulation electrodes, such as 200 µm Ag microwires (artifact size of ∼1mm), which are routinely used in functional MRI experiments.^[46,47]^

**Figure 7:**
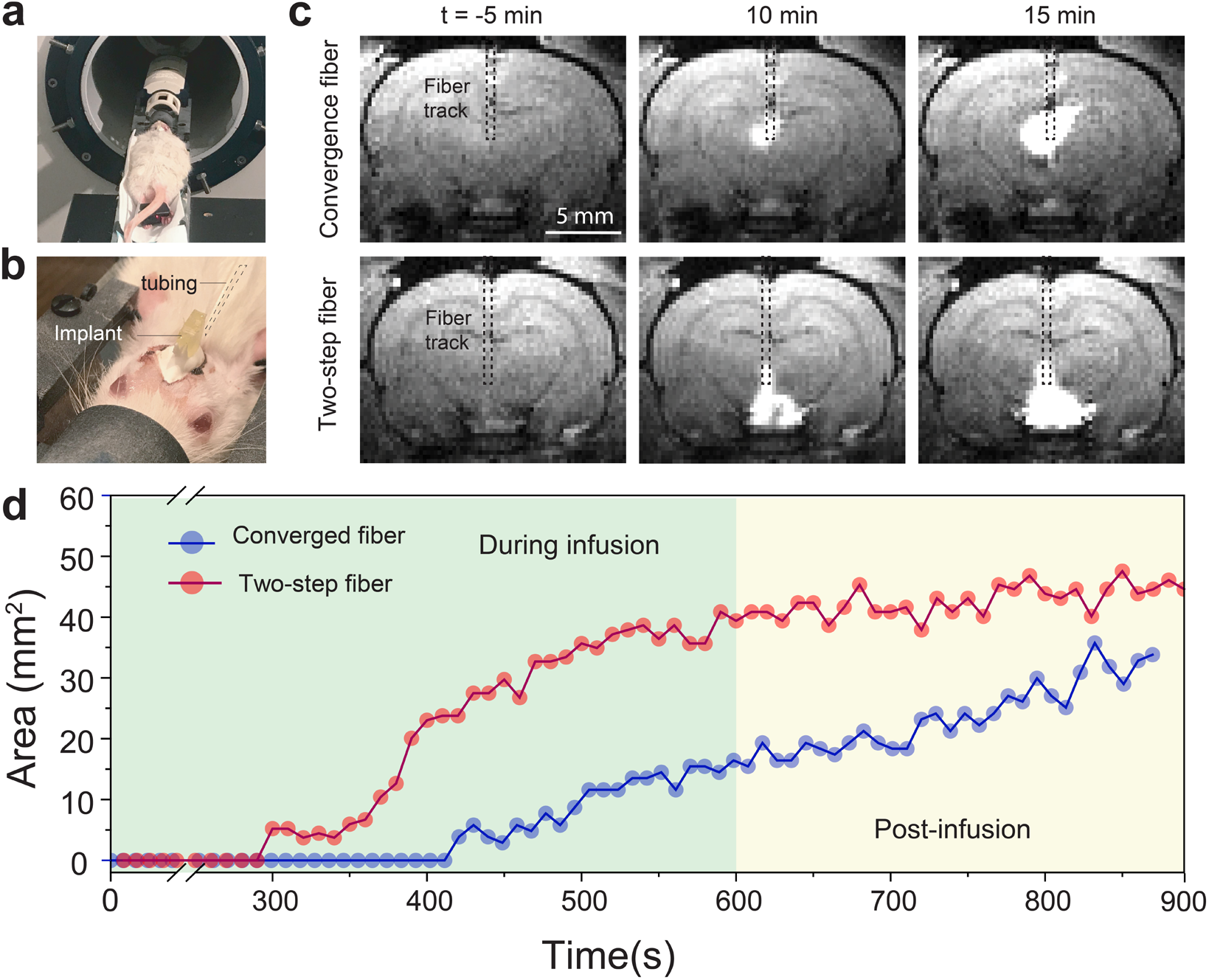
MRI in the presence of fiber-based probes. (a) Rat in MRI bore with surface coil placed over fiber implant. (b) Rat with fiber implant attached to infusion tubing. (c) MRI images of T1-weighted rapid acquisition with refocused echoes (RARE) pulse sequence pre (t = −5min), during (t = 10 min) and post (t = 15 min) injection of contrast agent through both fiber implants (convergence (top) and two-step (bottom) fiber). (d) Infusion profile showing contrast changing area over time as a measurement of infusion efficiency of the two fiber types over the course of a 20 min scan.

Although chemical neuromodulation methods are common in basic and translational research, the real-time visualization of neuromodulatory agents *in vivo* remains challenging.^[48]^ For example, imaging of the intracerebral fluid injection (as described above) is typically performed postmortem using standard histology techniques. This approach, however, is not suitable for tracking the delivery of fluid cargo in real time, nor can it ascertain the spatiotemporal kinetics of the infusion in intact animals.^[48]^ Here, we leverage the MRI compatibility of our multifunctional fibers to visualize in real-time microliter scale infusions of a Gd^3+^-based contrast agent ProHance (0.01 mM) in a deep subcortical region of the rat brain. We infuse this contrast agent at a rate of 20 nL/s over a 10 min period while MRI images are captured at a sampling rate of 0.1 Hz. **Figure 7c** shows T1-weighted rapid acquisition with refocused echoes (T1-RARE) images of implanted fibers of both types before, during, and following infusion. The bolus of the Gd^3+^ agent is clearly visible as an enhanced contrast around the fiber tip. As expected, the delivered fluid continues to diffuse out and away from the fiber tip following active infusion. **Figure 7d** illustrates the spatiotemporal kinetics profile of the contrast agent infusion by tracking the gradual increase of affected tissue area over time. **Videos S2** and **S3** show the complete infusion sessions in real time, while **Figure S15** and **Figure S16** depict the portfolio of snapshots captured every 10s from these videos. These experiments validate the MRI-compatibility of the multifunctional fibers for both anatomical and functional neuroimaging scans. They also motivate future studies aimed at coupling the local multimodal interrogation of neural circuits with global brain-wide activity mapping with MRI.

### Immunohistochemical evaluation of biocompatibility of the fiber probes

To assess the tissue response of the two-step fibers containing indium and converged multifunctional fibers containing tungsten, we performed immunohistochemical analysis in 50 µm thick coronal brain slices two and twelve weeks following implantations into the mPFC in mice. Similarly sized silica fibers (300 µm diameter, Thorlabs) were used for comparison given their prevalence in systems neuroscience studies (**Figure S17)**. Analysis of markers associated with the astrocytic response (glial fibrillary acidic protein, GFAP) and microglial activation (activated macrophage marker ionized calcium-binding adaptor molecule 1, Iba1)^[49,50]^ at two weeks following implantation revealed substantially lower tissue responses for the polymer fibers compared to silica fibers, which is consistent with prior reports of polymer based fibers.^[2,4]^ This trend persisted for microglial activation twelve weeks following implantation, while we observed a significantly lower astrocytic accumulation around the fiber-based probes as compared to silica (GFAP - P<0.01, **Figure S17**).

## Conclusion

Multimaterial fiber fabrication has enabled straightforward integration of neural recording, optogenetics, and drug delivery within miniature and flexible probes. However, previously thermomechanical compatibility at drawing conditions limited electrodes materials in fiber-based probes to either metals incompatible with low-loss waveguides, or polymer composites with high impedance. Through the use of an iterative thermal drawing with low T_m_ metal indium and convergence drawing of tungsten microwires, we produced an array of flexible multifunctional neural probes with low-impedance metal electrodes, low-loss polymer optical waveguides, and microfluidic channels. Notably, these fabrication advances led to a streamlined back-end interfacing of multifunctional probes.

We demonstrated the utility of fibers drawn using both approaches *in vivo* through chronic recording of spontaneous and optically-evoked electrophysiological activity in mice. Furthermore, we showed that these fiber-based probes could be combined with a mechanical microdrive to interrogate multiple clusters of neurons along the dorso-ventral axis in awake, freely moving mice. Finally, we found that both types of fibers were MRI-compatible and could be coupled with anatomical and functional brain-wide imaging while causing negligible tissue distortion artifacts.

This study expands the palette of available materials for fiber-based neural interfaces. By highlighting a streamlined connectorization process, the coupling of fiber-probes with mechanical microdrives, and their compatibility with MRI, this work aims to facilitate dissemination and widespread adoption of fiber-based multifunctional neural probes within the neural engineering and neuroscience communities.

## Materials and Methods

### Fabrication of the Two-step neural probes

The two-step fibers implants were fabricated using a combination of polycarbonate films (PC film, 50 μm thick, Ajedium; PC rod, McMaster-Carr) and cyclic-olefin-copolyer (COC 6015, 0.003” thick; TOPAS) for the waveguide and the microfluidic channel, and indium (In; 99.999% pure indium shot, Indium Corporation) as the electrode material. Two iterations of the thermal drawing process were used to obtain the final device. During the first iteration, optical waveguide, insulated electrode and microfluidic channel were individually fabricated and drawn. To fabricate the optical waveguide, 0.003” COC films were rolled around a 1” polycarbonate rod PC until the assembly reached 1.25” thickness. The microfluidic channel was fabricated by drilling a 1/2” hole through an 8” long, 1” thick PC rod. 0.003” COC was then wrapped onto the rod until it reached 1.25”. To fabricate the insulated electrode, a 3/8” hole was drilled into a 1” thick, 8” long PC rod, which was also wrapped in 25 µm COC sheets to reach 1.5” in thickness. All three preforms were consolidated at 175°C under vacuum for 40 min. The electrode channel was then filled with indium pellet, and all three preform were drawn at 275°C using a custom-built fiber drawing tower as described in previous work.^[2]^ The microfluidic channel was drawn with a draw-down ratio of 15 to obtain ∼2 mm fibers, the optical waveguide with a ratio of 30 and the electrodes a ratio of ∼40 to obtain ∼1 mm fibers. To fabricate the preform for the second iteration of the thermal drawing process. A COC/PC cylinder was fabricated by first rolling 1 layer of 0.001” fluorinated ethylene propylene (FEP) teflon (McMaster-Carr) around a 4.5mm steel mandrel (McMaster-Carr), followed by several sheets of COC sheets to reach a thickness of 5.25mm. It was then wrapped into PC films to reach 1” in thickness, this sacrificial cladding is necessary to enable stable drawing conditions. The assembly was consolidated under vacuum at 175°C for 40 min. The FEP wrapped steel mandrel was then removed, and optical, microfluidic and insulated electrode fiber were inserted. To eliminate air pockets between individual fiber strands, PC fibers were added, and the assembly was annealed in a vacuum oven at 110°C for 1 hr, before being drawn at 275°C with a draw down ratio of 35 to obtain the 2nd step fibers.

### Assembly of the two-step neural probes

Connectorization refers to the process of interfacing the individual modalities of neural probes (electrodes, microfluidic channel and optical waveguide) with input/output electric pins, inlet tube and optical ferrules. The fiber was first cut into 7 cm long sections, and the proximal end of each fiber was scored with a razor blade, snapped and pulled apart to expose the COC-wrapped first-step fibers. The COC layer was scored and pulled apart, releasing the individual first-step fibers. Once exposed, each individual fiber was connected to the corresponding hardware. The neural probe was placed on a 3D-printed box (microdrive shuttle), and individual fibers were connected to the corresponding hardware. Individual electrodes were connected and fixed to the male pin connectors (Digi-Key, #1212-1328-ND) using silver epoxy (Epo-Tek, #H20E). An insulated stainless-steel ground/reference wire (A-M systems, #790500) was soldered to one of the pins. EVA tube (0.03” ID, 0.09” OD, McMaster #18883T2) was placed around the microfluidic channel, and affixed to the fiber using UV epoxy (NO68; Norland). The optical waveguide was placed into a 10.5 mm long, 2.5 mm diameter ceramic ferrule, with a 270 µm bore size (Thorlabs, #CF270) and affixed it using optical epoxy (NO68; Norland). The silver epoxy was cured by placing the assembly onto a hot plate at 80 °C for 2 hr. UV-epoxy was added onto the box cavity to submerge the assembly for additional mechanical stability and electrical insulation. The ferrule was then polished using a Thorlabs fiber polishing kit, and the distal end of the implant was immersed into dichloromethane (Sigma, #270997) for 4 min to remove the sacrificial PC layer.

### Fabrication of the converged neural probes

Multifunctional fibers using the convergence approach were fabricated as follows. A macroscopic preform was assembled by rolling multiple layers of COC thin films (COC 6015, 0.003” thick; TOPAS) around a 1/8” PC rod that defined the central PC/COC waveguide scaffold. Subsequently an additional layer of PC was added by rolling PC films (film, 50 μm thick, Ajedium; PC rod, McMaster-Carr) until a diameter of ∼6.2 mm was reached. At this step the rolled preform was consolidated in a vaccum oven at 185 °C for 30 min. Next, six square slots of 1mm each were machined in a symmetrical pattern in the outermost PC layer which overall defined the four convergence channels and two microfluidic channels. Teflon spacers were inserted into these channels and additional PC sheets were rolled until a diameter of ∼9 mm was reached. Finally, alternate layers of COC films (0.003” thickness) and PC films (50 μm thickness) were rolled that defined the stopping layer and the cladding layer respectively. The assembled preform was again consolidated in a vacuum oven at 185 °C for 40 min. All the fiber drawing processes were conducted using a custom-built fiber drawing tower. The fibers were drawn by placing the preform in a three-zone heating furnace, where the top, middle, and bottom zones were heated to 150 °C, 285 °C, and 110 °C, respectively. The preform was fed into the furnace at a rate of 1 mm/min and drawn at a speed of 1600 mm/min, which resulted in a draw-down ratio of 40. Four tungsten microwire spools (25 µm diameter, 99.95%, Goodfellow) were continuously fed into the preform during the draw that defined the convergence process.

Tetrode-type electrophysiology fibers were fabricated as follows. A macroscopic preform was assembled by machining four grooves (1 mm) along the length of mm PC rod (1/8”) that defined the convergence channels. Teflon spacers were inserted into the hollow channels and layers of PC films (50 μm thin) were rolled until a final diameter of ∼4 mm was reached. The assembled preform was consolidated in a vacuum oven at 185°C for 20 min and drawn into meters long fibers using identical draw parameters as described above.

### Assembly of the converged neural probes

In a typical connectorization protocol for multifunctional fibers, the sacrificial cladding (PC) and the stopping layer (COC) were dissolved by subsequently submerging the fibers in dichloromethane (DCM) and cyclohexane for 5 and 10 min respectively. An intermediate wash cycle with IPA was performed before switching between the solvents. After etching the cladding layers, the fibers were dipped in DCM for 5 min to expose the individual metal microwires and PC/COC waveguide. The microwires were soldered to header pins (Digi-Key, #1212-1328-ND) and the waveguide was affixed into a ceramic ferrule (Thorlabs, #CF270) using UV epoxy (NO68; Norland) and subsequently polished with Thorlabs polishing kit. The microfluidic channels in the fiber were connected to an external EVA tubing (0.03” ID, 0.09” OD, McMaster #18883T2) by mechanically etching the polymer cladding on the fiber with a razor blade and then threading through the fiber into the tubing to form a T-junction. The assembly was made watertight using copious amounts of UV epoxy. The device assembly was finished off by soldering a ground screw to a reference wire on of the header pins and securing all the I/O hardware (pins, ferrules, tubes) inside a 3D printed Microdrive shuttle with copious amounts of UV epoxy. The tetrode type fibers were connectorized in a series of almost identical steps. To begin with, the fibers were etched in DCM for 5 min to expose all microwires. These electrodes were soldered to header pins. A reference wire connected to a ground screw was attached to one of the header pins and the whole backend assembly was supported inside a microdrive shuttle using copious amounts of UV curable epoxy.

### Characterization of two-step and converged neural probes

To image the neural implants, we embedded the fiber samples in an epoxy matrix (EMSFix Epoxy Resin) and grinded the sample with a series of sandpapers with decreasing grain sizes (Struers Tegrapol) before polishing with a 0.3 µm alumina particle solution (Buehler AutoMet 250 Polisher). Images were acquired using a metallurgical microscope. To highlight the functional part of the fiber and mask the sacrificial cladding, morphological image processing was performed by segmenting foreground and background pixels using a region-growing algorithm in Mathematica 12.0. Bending stiffness of the fibers were measured using a dynamical mechanical analyzer (Q800, TA Instruments). We used a single cantilever mode with 20 μm deformation within the frequency range of 0.1-10 Hz. Electrical impedance spectroscopy of fiber integrated electrodes was measured using a LCR meter (HP4284A, Agilent Technologies) with a sinusoidal voltage input (10 mV, 20 Hz-10 kHz). Microfluidic capability of the channel was characterized by implanting the fiber tip into a phantom brain (0.6 % wt/vol agarose in distilled H_2_O). The microfluidic inlet was connected to a syringe pump (WPI Micro4, UMC4) and controller (WPImUMP3) via a Hamilton syringe and EVA tubing (0.03”ID, 0.09”OD, McMaster #18883T2). Tryptan blue (3 µl) was injected at a speed of 750 nl/min, and images were captured at t=0, t=3 and t=5 following the start of the injection using a phone camera. To further characterize the microfluidic capabilities of the implants, 9 µl of water was injected at infusion speeds of 1, 10, 20, 50, 80 and 100 nl/s through the microfluidic channel, measuring injection output by weight. Output injection rate was calculated by dividing the injection output by the time required to inject it. Injection efficiency was computed as the ratio of input and output injection rate. To image the light emission profile of polymer waveguides, 5 cm long fiber probes were connected to a diode-pumped solid state (DPSS) laser (50 mW maximum output, wavelength λ=473 nm, Laserglow) via optical cable and a zirconia ferrule. The fiber tip was placed into a fluorescein solution (0.01mM), light was delivered through the fiber and images were captured. Transmission losses for waveguides were measured by varying the fiber lengths from 1 to10 cm using the cut-back method. The absolute output power was measured using a digital power meter (Thorlabs, PM100D) with a photodiode sensor (S121C). All the electrochemical measurements were performed with a potentiostat (Solartron, SI 1280B) at room temperature. The bulk indium rods (In; 99.999% pure indium shot, Indium Corporation) or two-step indium fibers were utilized as working electrodes. Pt wire (MW-1032, BAsi) and Ag/AgCl electrode (BASi, 3 M NaCl) were used as the counter and reference electrodes, respectively. Phosphate buffer saline (PBS, Fisher Scientific) was used as the electrolyte, and cyclic voltammetry curves for the indium working electrodes were obtained at the scan rate of 20mV/s. Potentiostatic electrochemical impedance spectroscopy (PEIS) was performed at the open circuit potential (OCV) with a Biologic VMP3 potentiostat on first-step indium fibers in a PBS electrolyte. The working electrode was held at open circuit for 2 sec, followed by scanning frequencies from 1 MHz to 10 Hz at six points per decade. An amplitude of 5 mV was used, and an average of 10 points was performed at each frequency. A leak-free Ag/AgCl reference electrode (Innovative Instruments) and a platinum foil counter electrode (99.99% trace metals basis, Beantown Chemical) were used. Equivalent circuit fitting was performed with Z Fit, an EIS fitting tool in EC-Lab with 5,000 iterations using a randomize + simplex algorithm. A simplified Randles circuit model, consisting of a solution resistance in series with a parallel combination of a charge-transfer resistance and a double-layer capacitance, was used to fit the data; the double-layer capacitance was modeled both as an ideal capacitor and as a constant phase element, although the ideal-capacitor model was chosen for its simplicity and lower uncertainty in its fitted parameters. The finite element modelling software, Abaqus, is used for the simulation of the bending of the different fibers as well as the fiber displacement in moving tissue. The element used for the materials in the fiber, such as the polymer, silica, and metal wires, is C3D8H with a HEX element shape. The element used for the tissue is C3D10H with a TET element shape. For the bending fiber simulation, the fiber is held fixed at one end and is displaced with a fixed distance at the other end. The amount of displacement is 0.1 mm. For the simulation for the fiber in the brain tissue, the tissue at its bottom is displaced sidewards with a distance of 3.5 um, while the exposed end of the fiber, which is not in the tissue, is fixed.

### Microdrive fabrication and characterization

The microdrive body, shuttle and cover were designed on Solidworks (Dassault System) and 3D printed on a Form 2+ (Formlabs) using a clear resin (Formlabs, RS-F2-GPCL-04), at a 25 μm resolution. Following 3D printing, each element was cleaned using the FormWash for 30 min, then cured for 180min into the FormCure. Due to the imprecision of 3d printing at this scale, screw channels onto the body and the shuttle were widened to the designed 0.5mm using a handheld drill. Following connectorization of the fiber onto the shuttle. Microdrive body, shuttle carrying the fiber, and custom-made screw were assembled. The drive body and shuttle were assembled by placing the shuttle into the guide channel, and using a custom screw described in Voigts et al.^[41]^ The shuttle was then place to its uppermost position, and linear motion of the screw was prevented by filling the retaining pocked with light-hardened dental cement (Vivadent Tetric EvoFlow). The screw was then slightly turned counterclockwise to delaminate the screw from the cement that now acts as a thrust bearing.

Quantitative microdrive characterization was performed by fixing the stationary part of the device to a glass slide. A rectangular cuvette filled with 0.6% agarose was fixed to another glass slide. The fiber was introduced into the agarose gel, and the slides were both fixed into place under a light microscope. The microdrive was then advanced one turn at a time, and microscope images were acquired after each turn. The distance traveled by each turn was analyzed by measuring the number of pixels between the end of the fiber and a reference point on the cuvette. Pixels were converted to micrometers by measuring the diameter of the fiber. To illustrate the performance of the microdrive, the device was fixed to the top of a glass vial filled with 0.6% agarose. Three injections of 500 nl of Evans Blue (2% in PBS) were delivered at a rate of 100 nl/min at depths of 0, 1 and ∼2 mm (7 and 14 turns).

#### Surgical procedures

All animal procedures were approved by the MIT Committee on Animal Care and performed in accordance with the IACUC protocol 0121-002-24 (Implantation of Neural Recording and Stimulation Devices into Adult Mice and Rats). WT mice (Jacskon, 000064) aged 8 weeks, were used for the study, and housed in a normal 12h light/dark cycle and fed a standard rodent chow and water diet *ad libitum*. All surgeries were performed under aseptic conditions. Mice were anesthetized with 1-2% isoflurane and placed on a heat pad in a stereotaxic head frame (Kopf Instrument), and were injected subcutaneously with slow-release buprenorphine (ZooPharm, 1.0 mg/kg). Following a midline incision along the scalp, the skull was repositioned by aligning and leveling Lambda and Bregma landmarks with respect to the Mouse Brain Atlas. Adeno associated viruses serotype 5 (AAV5) carrying CaMKIIα::hChR2(H134R)-eYFP and CaMKIIα::EYFP plasmids were purchased from University of North Carolina Vector Core at concentrations of 3 × 10^12^ particles/ml and 2 × 10^12^ particles/ml, respectively. Implantation and injection coordinates were established following the mouse brain atlas. A craniotomy was performed using a rotary tool (Dremel Micro 8050) and a carbon steel burr (Heisinger, 19007-05). The animal was then either injected with virus or implanted with the neural probes. Virus injection was performed using a standard NanoFil Syringe and UMP3 Syringe pump (World Precision Instruments), 0.5 μl of virus was injected using a at a rate of 100nl/min. Prior insertion, implants were flushed with sterile PBS, fastened to the stereotaxic cannula holder, and were connected to a RZ5D (TDT) recording system using a PZ2-32 head-stage with a 16-channel zero insertion force-clip headstage. Probes were lowered into the medial prefrontal cortex (mPFC; ML = 0.4mm; AP = +1.7mm; DV= 2.5mm). Following insertion, a stainless steel screw (McMaster-Carr, # 92196A051) was inserted into the skull above the cerebellum as a ground and reference. The device was then cemented to the skull using 3 layers C&B-Metabond adhesive acrylic (Parkell) then dental cement (Jet Set-4) to cover both the screw and the base of the device. The incision was then closed with 5-0 suture in a simple interrupted pattern, and the mouse was subcutaneously injected with carprofen (5mg/kg) and sterile Ringer’s solution (0.6 mL) prior to be returned to their home cage, partially placed on a heat pad. Post-implantation, the mice were provided food and water *ad libitum*, were monitored for 3 days for signs of overall health, and were given carprofen injection (0.6 mL, 0.25mg/ml in sterile Ringer’s solution) as necessary.

### In vivo electrophysiology

Neural implants were attached to a PZ2-32 head-stage with a 16-channel zero insertion force-clip connector which was in turn connected to a RZ5D recording system. Electrophysiological signals were filtered in the frequency range 0.3 - 5 kHz and digitized at a sampling frequency of 50kHz. Subsequent signals processing and analysis was done with Matlab (Mathworks). Spiking activity was detected using threshold detection with a threshold set at 5 standards deviation from the mean of the signals, with a downtime of 2ms to reject double detections. lustering and classification of spikes were performed by principal component analysis and Gaussian Mixture Model clustering (with full and independent covariance matrices). We assessed the quality of the clustered data by calculating the L-ratio and the isolation distance of the classified clusters.^[33]^

During optogenetic stimulation, a 500mW 473nm diode-pumped solid-state laser (OEM Laser Systems) was coupled to the implant through a silica fiber (200μm core, Thorlabs #RJPFF2) using a rotary joint optical fiber and a ceramic mating sleeve. Waveform generation was delivered by the RZ5D acquisition system with a stimulation frequency as set at 10 or 100hz, 5-ms pulse width, 1s trial duration and 4 s intertrial interval. During the control experiment with violet (λ = 420 nm) and green (λ = 565 nm) light emitting devices (LEDs), the optical stimulation was generated using fiber-coupled LEDs (Thorlabs, M420F2, M565F3) driven by a LED driver (Thorlabs, LEDD1b, 1.2A current limit) and a waveform generator (Agilent 33500B).

During chemical delivery, the implants were connected to a Nanofil syringe (10 µl, World precision Instruments) using an EVA tube (0.03” ID, 0.09” OD, McMaster) and a Nanofil beveled needle (33G, WPI). 2 µl of Evans Blue (2% in sterile NaCl) was injected at a speed 100 nl/min through the implants’ microfluidic channels.

### In vivo MRI imaging

Adult male Sprague–Dawley rats (350–450 g) were purchased from Charles River Laboratories (Wilmington, MA). After arrival, animals were housed and maintained on a 12-hr light/dark cycle and permitted ad libitum access to food and water. All procedures were carried out in strict compliance with National Institutes of Health guidelines, under oversight of the Committee on Animal Care at the Massachusetts Institute of Technology. Animals were anesthetized with isoflurane (3% for induction, 2% for maintenance) and placed on a water heating pad from Braintree Scientific (Braintree, MA) to keep body temperature at 37 °C. Animals were then fixed in a stereotaxic frame, and topical lidocaine was applied before a 3 cm lateral incision extending from bregma to lambda was made, exposing the skull. Craniotomies (0.5 mm) were drilled unilaterally over the ventral tegmental area (VTA), 6.0 mm posterior and 0.5 mm lateral to bregma. Fiber implants (two step or convergence) were preloaded with a gadolinium-based contrast agent (ProHance, Bracco Diagnostics; 0.01mM), lowered to 8.0 mm below the surface of the skull through the craniotomy and held in place by the stereotactic arm. The implants were fixed to the skull with dental cement (Secure Resin Cement, Parkell). Animals were then transferred into a custom rat imaging cradle, fixed with ear bars and bite bar, maintained under 2% isoflurane anesthesia, and kept warm using a recirculating water heating pad for the duration of imaging.

MRI was performed on a Bruker 9.4 T BioSpec small animal scanner (Billerica, MA) using a transmit-only 70 cm inner diameter linear volume coil made by Bruker (Billerica, MA) and a receive-only 3×1 array-coil with openings through which the implant and connected tubing was led through to then be connected to an infusion pump. Scanner operation was controlled using the ParaVision 6.01 software from Bruker (Billerica, MA).

Multislice T1-weighted fast low-angle-shot MRI images with TE = 4 ms, TR = 200 ms, field of view (FOV) = 30 × 18 mm, in-plane resolution 200 × 200 μm, and four to six coronal slices with slice thickness = 1 mm and T2-weighted TurboRARE images with TE = 30 ms, TR = 1000 ms, Rare Factor 8, Matrix size 256 × 256, FOV = 25.6 × 25.6 and in-plane resolution of 100 × 100 µm were used to evaluate the location of the implants.

Rapid acquisition with refocused echoes (RARE) pulse sequences were used for functional scans with the following parameters: FOV = 30 × 18 mm, slice thickness = 1 mm, matrix size = 128 × 64, and RARE factor = 4 (with 5 ms echo spacing). Functional scans of 20 min length were performed during the infusion of contrast agent over the course of 10 min (20nl/s), starting 5 min after the start of the scan.

### Behavior assays

Locomotor tests were performed with virally transfected mice, in a 30 × 30 × 30 cm^3^ white acrylic chamber. Optical stimulation was generated by blue laser (λ = 473 nm) controlled by a waveform generator (Agilent). The laser was collimated and coupled into a silica patch cord with a rotary joint optical fiber and were attached to the implant using a ceramic mating sleeve. Prior to the start of the behavioral experiment, mice were restrained while the patch cord was attached to their implant, and were placed in the center of the chamber for 10 min to recover from handling stress. Each session consisted of three 3-min epochs, wherein baseline locomotor activity was assessed in the first and third epoch (OFF stimulation) and optical stimulation with 20hz frequency (5ms pulse width) was delivered during the second epoch (ON stimulation). The locomotor activity was tracked and analyzed using Ethovision software. The ability of mice to carry light-weight microdrive implants was measured using an Open-field assay in a 30 × 30 × 30 cm^3^ white acrylic chamber. Mice were acclimatized to the OFT chamber for 10 min before recording their activity. The OFT test session lasted for 10 min during which the overall distance travelled and locomotion velocity was tracked and analyzed using Ethovision software.

### Immunohistochemical evaluation of foreign body response

Animals were anesthetized with isoflurane, injected with fatal plus (100 mg/kg IP), and transcardially perfused with 50 mL of ice-cold phosphate buffered saline (PBS) followed by 50 mL of ice-cold 4% paraformaldehyde (PFA) in PBS. The devices were carefully explanted and the brains were removed and fixed in 4% PFA in PBS for 24 hr at 4 °C, then stored in PBS afterwards.

Coronal slices (50 μm thickness) were prepared using a vibratome (Leica VT1000S) and a razor blade (Electron Microscopy Sciences, 72002) in ice-cold PBS. The slices were then stored in PBS at 4 °C in the dark until staining.

Slices were permeabilized with 0.3% (vol/vol) Triton X-100 and blocked with 2.5% donkey serum in PBS for 30 min. Slices were incubated overnight at 4 °C in a solution of primary antibody and 2.5% donkey serum in PBS. Following incubation, slices were washed three times with PBS. The slices were then incubated with secondary antibody (Donkey anti-Goat Alexa Fluor 488, A11055, 1:1000, Thermofischer) for 2 hr in room temperature on a shaker followed by additional three washes with PBS. Slices were then incubated with DAPI (4’6-diamidino-2-phenylindole) (1:50,000) for another 20 min, and washed three times with PBS. Fluoromount-G® (SouthernBiotech) was used for mounting slices onto glass microscope slides. A laser scanning confocal microscope (Fluoview FV1000, Olympus) was used for imaging with 20× objectives. A laser scanning confocal microscope (Fluoview FV1000, Olympus) was used for imaging with 20× objective, with z-stack images across the slice thickness. Region of interest was chosen based on the fiber location, imaging the immune response in the fiber surrounding.

The following primary antibodies were used:

| Target | Antibody | Dilution |
| --- | --- | --- |
| Iba1 | Goat anti- Iba1 (ab107159, Abcam) | 1:500 |
| GFAP | Goat anti- GFAP (ab53554, Abcam) | 1:1000 |

### Statistical Analysis

All data is presented as mean ± S.E.M. Statistical significance was assessed using Matlab (Mathworks), by first confirming a normal distribution using a Lilliefors test, and then by 2-sided t-test, with significance threshold placed at *P<0.05, **P < 0.01, ***P < 0.001.

## Author Contributions

P.A., M-J.A., and A.S. designed the study. M-J.A., A.C., and M.K. drew the two-step indium fibers. A.S. and T.K. drew the converged fibers. Y.F. provided advice in fiber processing. M-J.A. and A.S. respectively connectorized the two-step fibers, and the converged fibers. A.S., and G.L characterized the mechanical properties of the fibers, while M-J.A. A.S., J.P., and N.C. characterized the electrical properties, and M-J.A. and A.S. characterized the optical and microfluidic properties. M-J.A. and I.G. characterized the microdrive. M-J.A. and A.S. performed in vivo experiments and behavior. A.T. and A.S. perfused the animals, M-J.A., sliced the brain, D.R. and A.S. performed staining and confocal imaging. A.S. and M.S. performed the MRI experiments. A.P.J. offered insights into MRI and ensured access to the imaging facility. All authors have contributed to writing of the manuscript.

The authors declare no conflict of interest.

This article contains supporting information online.

## Acknowledgments

This work was supported in part by National Institute of Neurological Disorders and Stroke (R01-NS115025-01A1, P.A.), National Science Foundation (NSF) Center for Materials Science and Engineering (DMR-1419807, P.A. and Y.F.), National Institute of Health (U01 NS107712, A.J), NSF Center for Neurotechnology (EEC-1028725, P.A.), and the McGovern Institute for Brain Research at MIT (P.A.). The authors are also grateful to Georgios Varnavides for his help with image processing, Ayse Guvenilir for her help with fiber connectorization, and to Karthish Manthiram for assistance with equipment. A.T. thanks the Paul and Daisy Soros Fellowship and the NSF Graduate Research Fellowship for funding support. M.J.A. is a recipient of the Friends of McGovern Graduate Fellowship. A.S thanks the Lore Harp McGovern Fellowship for funding support. G.L., T. K. and Y.F. were supported by the MIT MRSEC through the MRSEC Program of the National Science Foundation under award number DMR-1419807, the US Army Research Laboratory and the US Army Research Office through the Institute for Soldier Nanotechnologies, under contract number W911NF-13-D-000, the MIT Sea Grant under the contract number NA18OAR4170105, and the Defense Threat Reduction Agency through the Department of Defense under the contract number SA21-03.

## Notes

### Competing Interest Statement

The authors have declared no competing interest.

