## Supplementary Figures and Videos for "Customizing Multifunctional Neural Interfaces through Thermal Drawing Process"

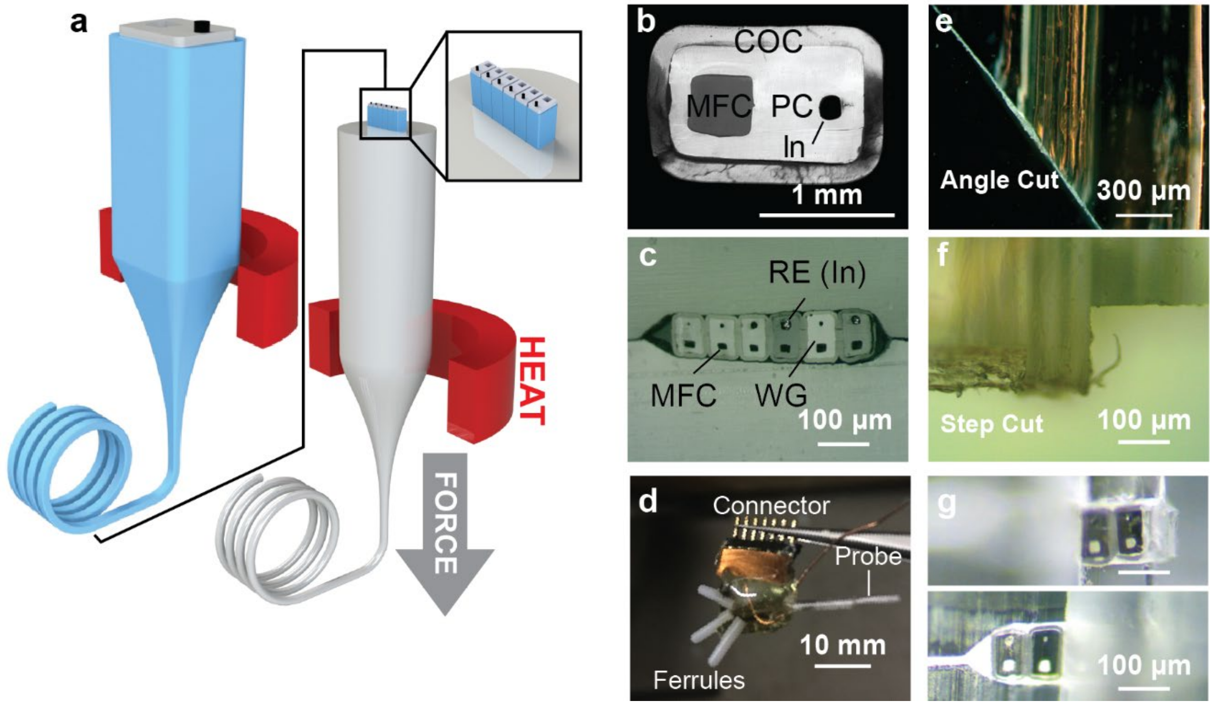


**Supplementary Fig. 1: Iterative thermal drawing for depth-specific neural probes**. Using two-step thermal drawing process (a) a preform with a PC core and a COC cladding, a microfluidic channel and an indium electrode is drawn into mm-scale fiber (b). These are then stacked into another PC rod and drawn into microscale fiber (c). (d) illustrate a connectorized device. Each repeating unit featuring an optical waveguide, a microfluidic channel and a recording electrode. An angle (e) or a step-cut (f, g) of this device enable depth-specific neuromodulation.

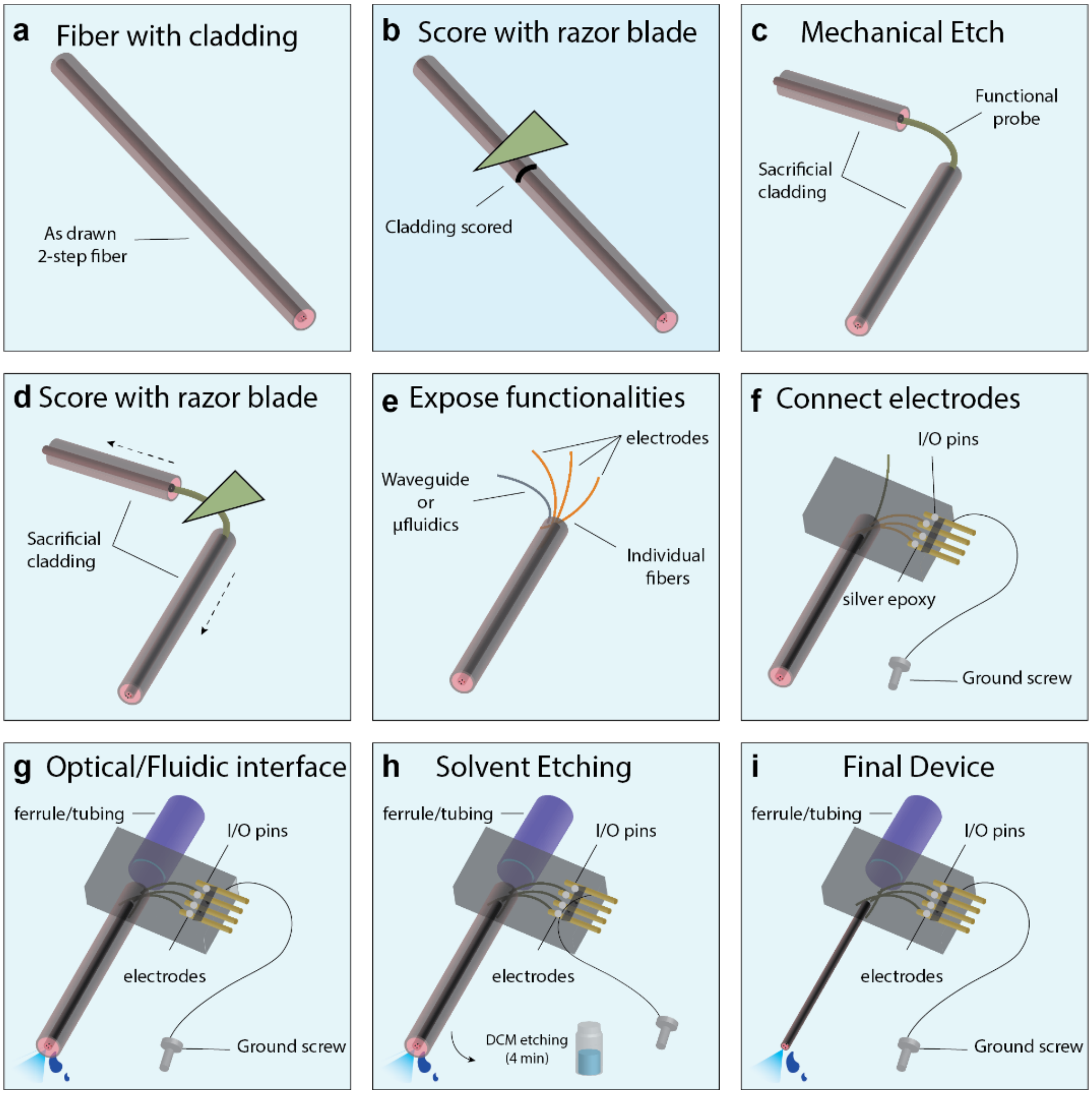


**Supplementary Fig. 2: two-step TDP enable rapid connectorization.** (a-i) Connectorization procedure for probes fabricated using the two-step thermal drawing (a) The probe is drawn with a protective sacrificial cladding. (b) A razor blade is used to score the surface of the sacrificial cladding. (c) Gently bending and pulling the fiber frees the neural probe from its cladding. (d) Scoring and pulling of the outer COC layer of the probe allows for the individual components to split apart (e). (f) The individual electrodes can then be connected to pin headers using silver epoxy (f), while the optical wavefuide and microfluidic channel can be connected using UV epoxy to an optical ferrule and fluidic tubing, respectively (g). The distal end of the probe is inserted into dichloromethane (DMC) for 4min (h) to remove the sacrificial cladding and expose the functional fiber.

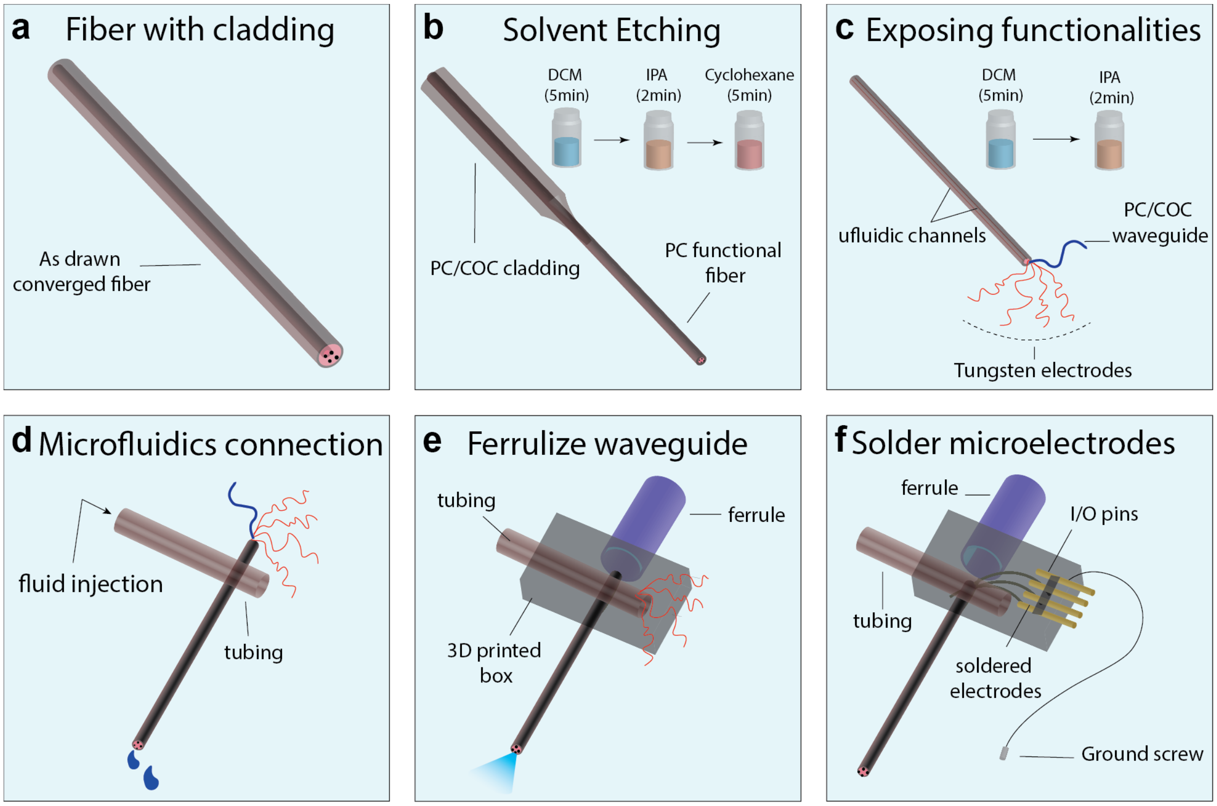


**Supplementary Fig. 3: Convergence TDP enable rapid connectorization.** (a-f) Connectorization procedure for probes fabricated using the convergence TDP (a) The fiber is drawn with a protective sacrificial cladding. (b) The sacrificial cladding of PC and COC are etched using a simple solvent etching process that involves alternate cycles of DCM (5min), IPA(2min) and cyclohexane (5min) wash (c) The embedded functionalities of microwire electrodes and PC/COC waveguide are exposed by solvent etching using alternate wash cycles of DCM (5min) and IPA(2min). (d) The microfluidic channel is connected to external tubing using a T-connection. (e) The exposed waveguide from step (c) is coupled to a zirconia ferrule and subsequently polished (f). Finally, the exposed microwire electrodes from step (c) are soldered to header pins along with a ground screw bearing reference wire. All backend I/O interfaces are encapsulated in a 3D printed shuttle drive using UV curing epoxy in (e,f).

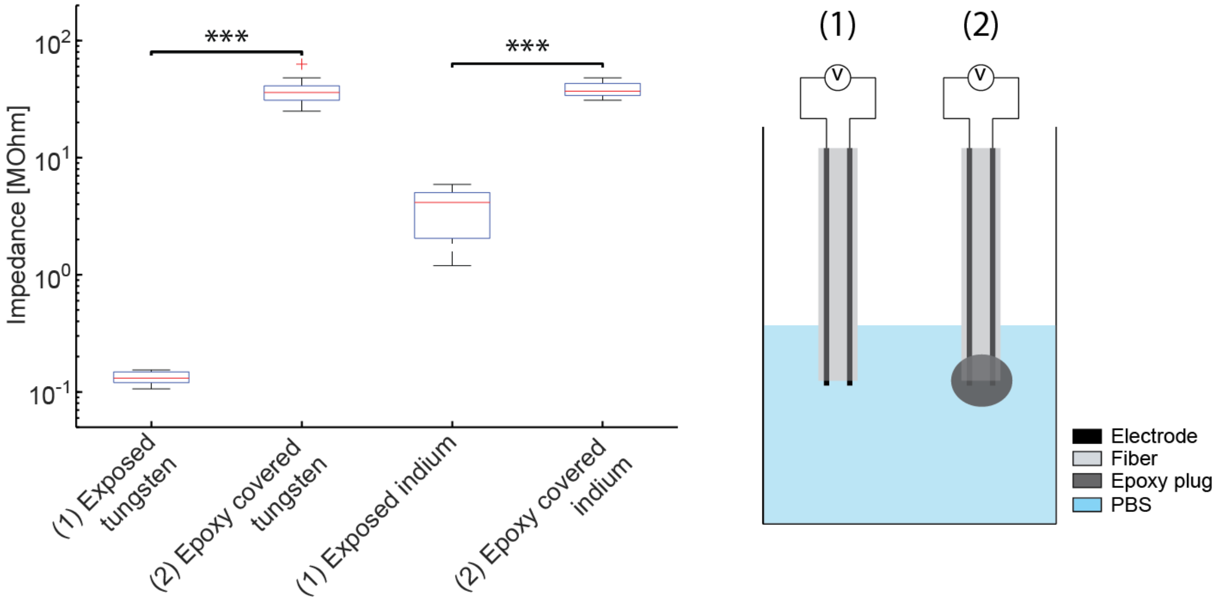


**Supplementary Fig. 4: Impedance across electrodes before and after covering the fiber tip with epoxy.** Impedances at 1kHz across electrodes within the two-step fiber (indium) and converged fiber (tungsten). (1) Impedance measurement across exposed electrodes at the fiber tip in PBS, (2) open-circuit measurement of the epoxy-insulated fiber tip in PBS. (N=10 samples)

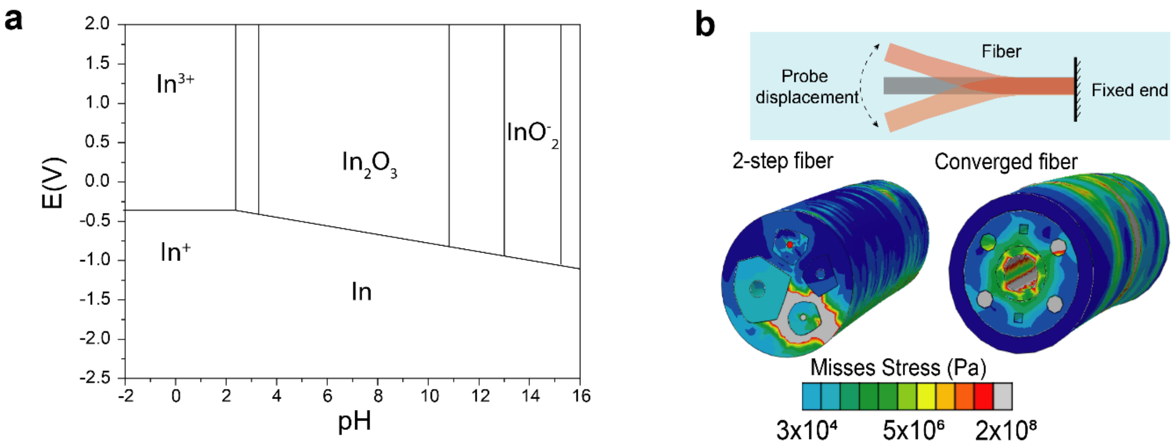


**Supplementary Fig. 5**: (a) Pourbaix (E vs. pH diagram) for indium in water at 25 °C. This diagram was reproduced from ref. [51]; (b) Misses stress profile under bending deformation computed using finite element models for two-step (left) and converged fibers (right), respectively.

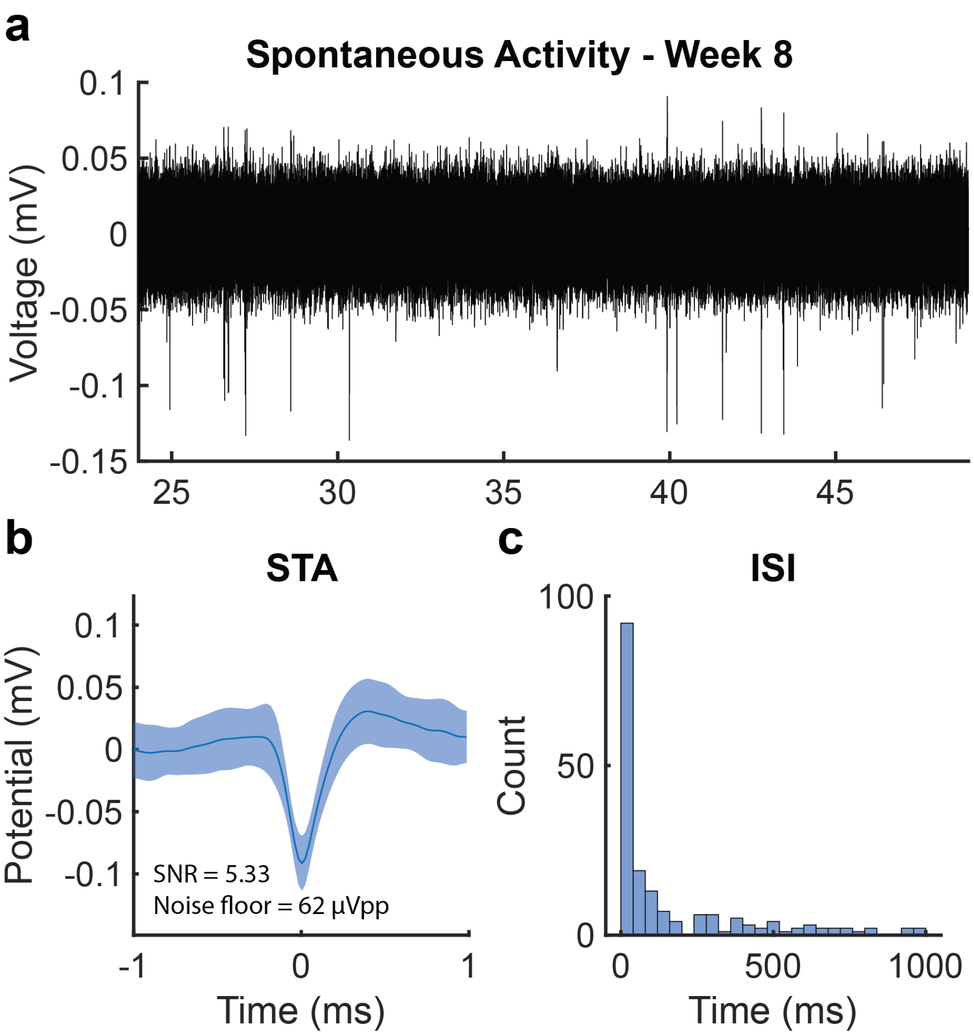


**Supplementary Fig. 6**: Two-step indium fiber records single unit activity 8 weeks post-implantation.

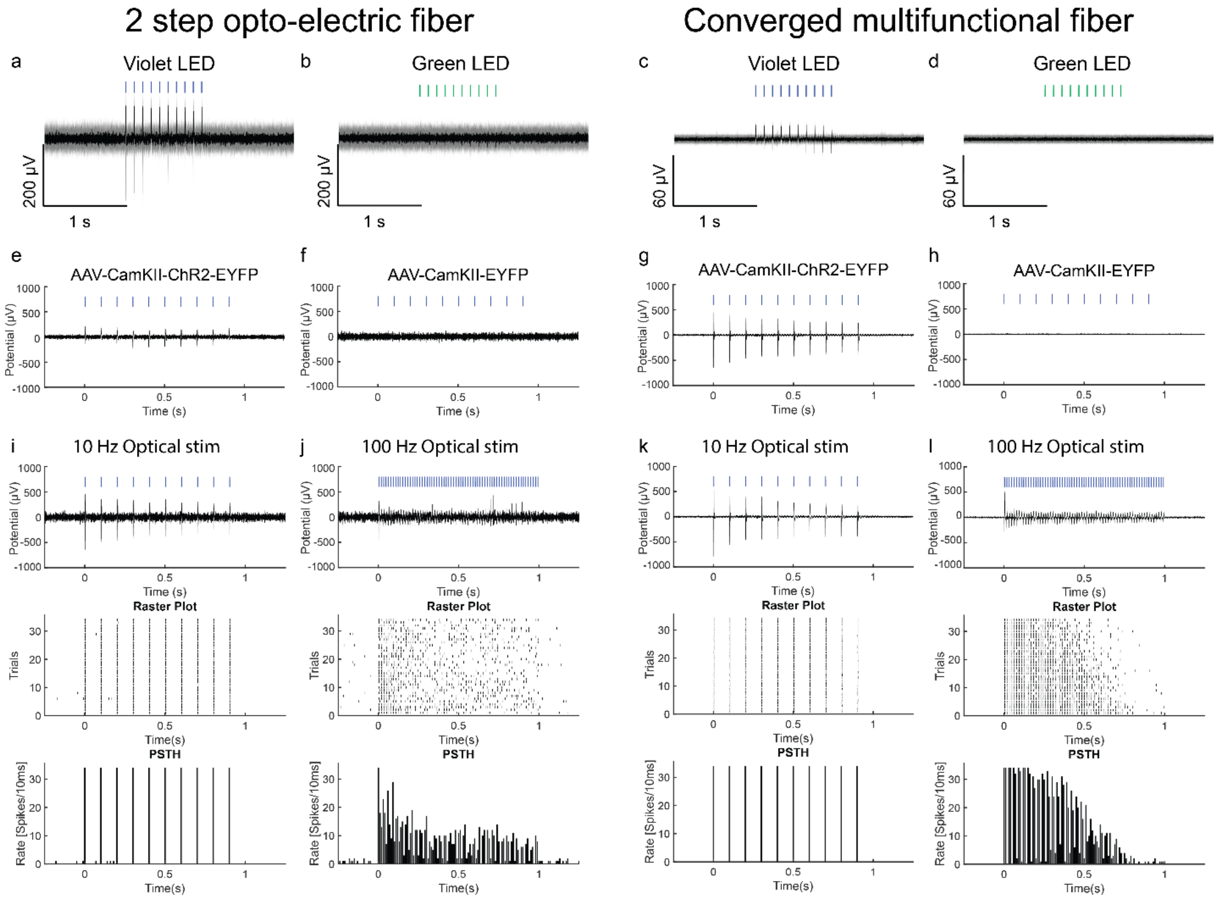


**Supplementary Fig. 7:** Controls experiment for optically evoked activity using the two-step and converged fiber implanted in the mPFC. (a-d) Comparison of electrophysiological response to green LED (565 nm, 10 ms, 10 Hz, 4 mW/mm$^2$) and violet LED (420 nm, 10 ms, 10 Hz, 4 mW/mm$^2$) light, for the two-step optoelectric and the converge multifunctional fiber (blue markers indicate laser onset). Only blue irradiation elicits electrophysiological activity. (e-h) Optically-evoked electrophysiological activity in mice transfected with AAV5-CaMKIIα::ChR2-EYFP (or control virus AAV5-CaMKIIα::EYFP). Neuronal activity is observed in mice transfected with ChR2, and not in control. (i,j) Comparison of optically evoked activity (473 nm, 10 ms, 10 mW/mm$^2$) at 10Hz and 100 Hz optical stimulation. Steady and correlated activity is observed at 10 Hz (i,k), as seen on the neural response (top), its associated raster plot (middle) and peri-stimulus time histogram (bottom). In comparison, 100 Hz stimuli elicited decaying and uncorrelated activity (j,l)

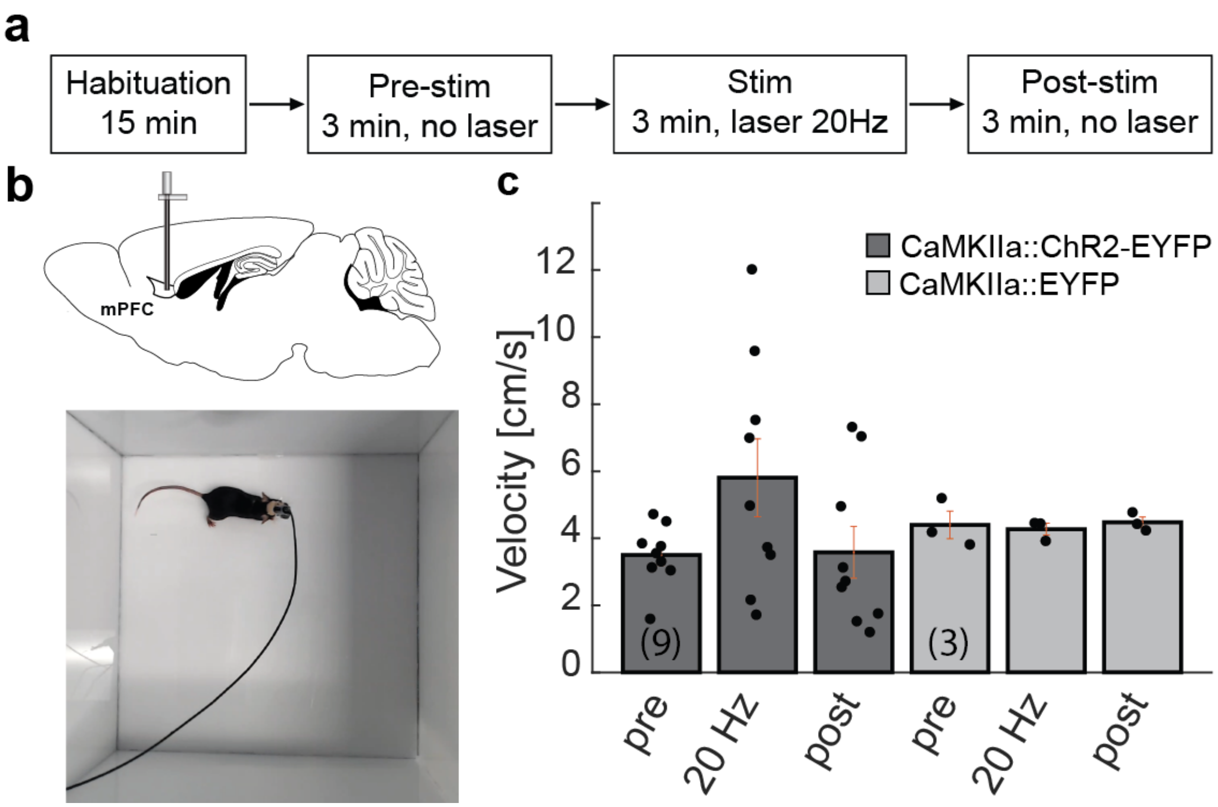


**Supplementary Fig. 8 – Behavioral assessment of mice implanted with converged and two-step fibers during optical stimulation.** (a) Schematic depicting the Open Field Test paradigm. Animal that were implanted with the multifunctional converged or the two-step opto-electric fiber (b, top) were first left habituated to the behavioral chamber for 15 min (b, bottom), then recorded over a 9-min experiment, consisting of 3-min OFF epoch, 3-min ON epoch during which they were exposed to 20 Hz stimulation (5-ms pulse width), 3-min OFF epoch. (c) Average velocity recorded for WT mice transfected with AAV5-CaMKIIα::ChR2-EYFP (or control virus AAV5-CaMKIIα::EYFP), then implanted with two-step and converged neural probe in mPFC.

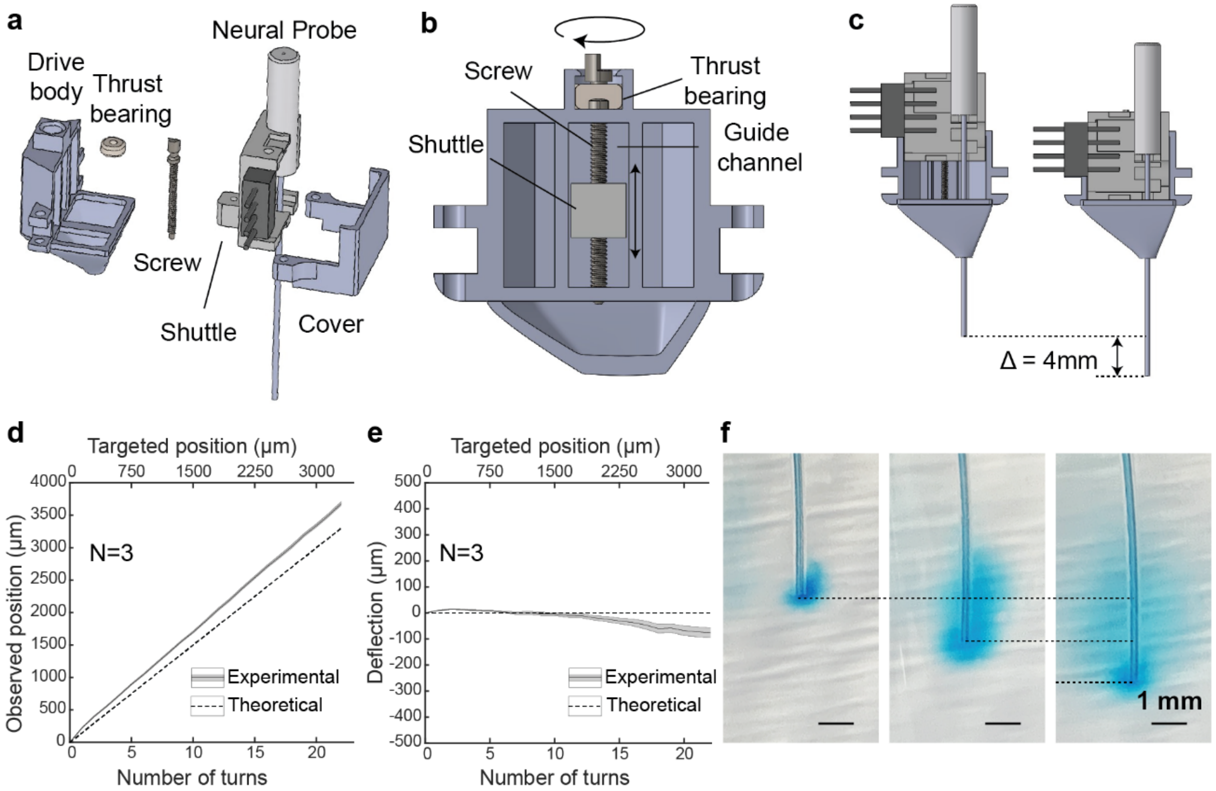


**Supplementary Fig. 9:** Fabrication and characterization of a microdrive for post-implantation positioning of multifunctional neural probes. (a) Exploded view of the microdrive assembly composed of a 3d-printed drive body, a dental cement thrust bearing, a custom-made screw, a neural probe mounted on a shuttle, and a 3d-printed cover. (b) Schematic of the cross section of the microdrive illustrating the screw and shuttle system. The shuttle moves linearly via actuation of the screw, the screw rotate inside a guide channel and is held linearly by the light-cured cement. (c) Schematic of the shuttle in the upper drive and lower drive position, illustrating a range of 4mm. (d) Measurement of the probe depth relative to the number of screw turns relative to top position. (e) Lateral deflection of the probes relative to the number of screw turns. (N=3) (f) Injection of Evans Blue dye (2%; 500nl over 5 min) into a phantom brain (0.6% agarose gel) with the shuttle at 3 different position (0, -1mm, and -2mm).

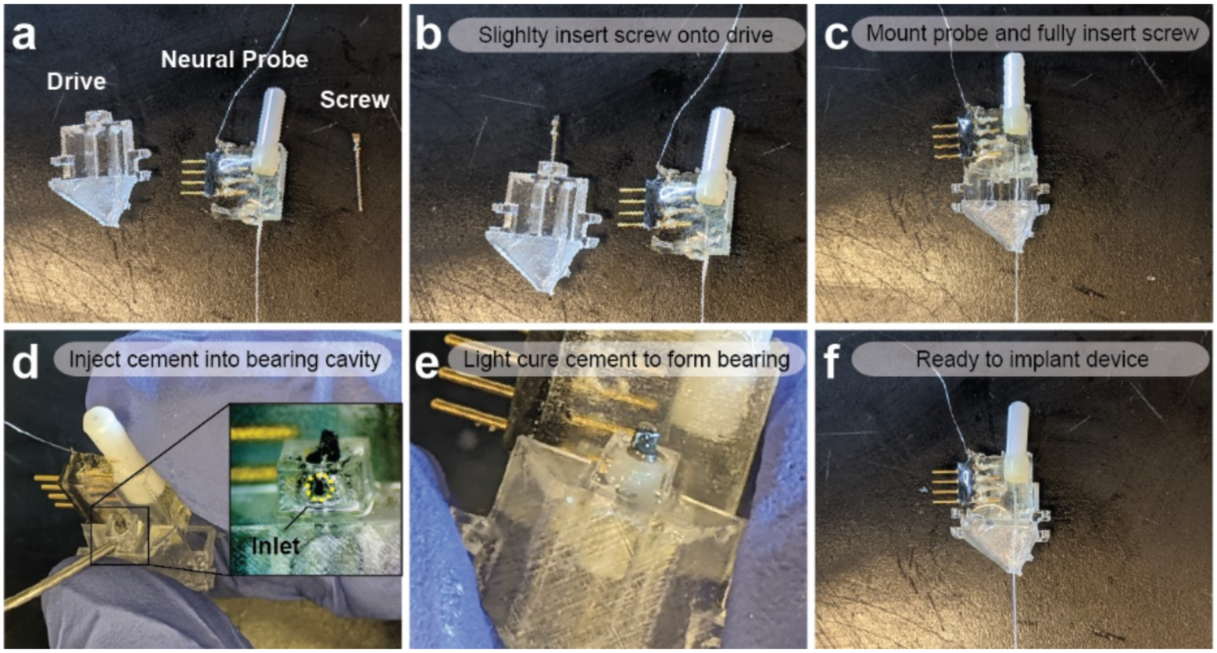


**Supplementary Fig. 10: Assembly procedure of the neural probe onto the microdrive.** (a) The microdrive assembly is composed of the 3d printed drive, the neural probe and a custom-made screw. (b) First, insert the screw into the screw slot on the top of the drive. (c) Place the neural probe shuttle near the bottom position in the guide channel, with the screw hole aligned vertically, and insert the screw into the shuttle hole. (d) Inject light-cured cement into the bearing cavity at the top of the drive, and (e) cure-it around the screw collar to form a thrust bearing that prevents linear motion of the screw. (f) Rotate the screw slightly to separate the screw from the bearing, the device is ready to be implanted.

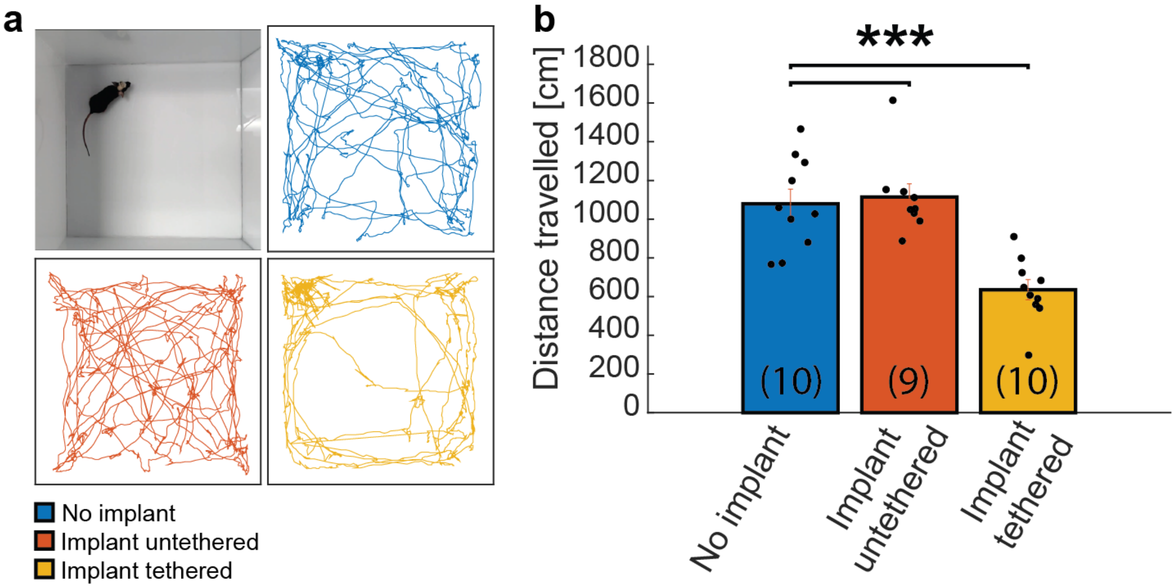


**Supplementary Fig. 11: Comparison of locomotor activity with and without the microdrive implant.** (a) Picture of the open field test (top, left), and representative motion tracks of mice during a 3-min exploration session (top right, bottom left and right). (b) Total distance travelled over a 3-min exploration session indicates that the mice are not hindered by the implant, but exhibit decreased activity when the microdrive is tethered to external hardware (top, right). *** P<0.001.

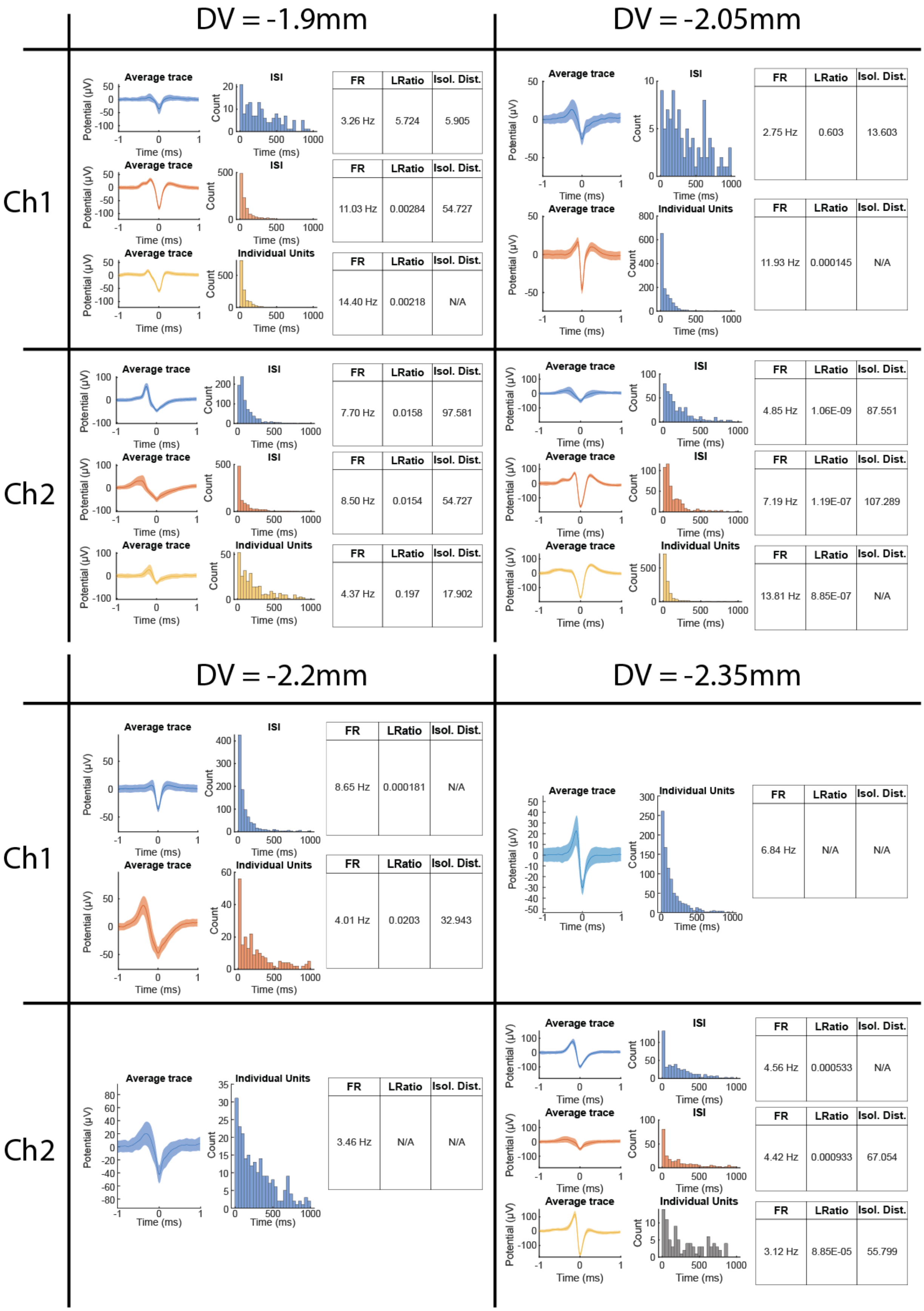
DV = -1.9mm
DV = -2.05mm
Ch1
Ch2
Average trace
ISI
Individual Units
Potential (µV)
Count
Time (ms)
FR | LRatio | Isol. Dist.
3.26 Hz | 5.724 | 5.905
11.03 Hz | 0.00284 | 54.727
14.40 Hz | 0.00218 | N/A
2.75 Hz | 0.603 | 13.603
11.93 Hz | 0.000145 | N/A
7.70 Hz | 0.0158 | 97.581
8.50 Hz | 0.0154 | 54.727
4.37 Hz | 0.197 | 17.902
4.85 Hz | 1.06E-09 | 87.551
7.19 Hz | 1.19E-07 | 107.289
13.81 Hz | 8.85E-07 | N/A
DV = -2.2mm
DV = -2.35mm
Ch1
Ch2
8.65 Hz | 0.000181 | N/A
4.01 Hz | 0.0203 | 32.943
6.84 Hz | N/A | N/A
3.46 Hz | N/A | N/A
4.56 Hz | 0.000533 | N/A
4.42 Hz | 0.000933 | 67.054
3.12 Hz | 8.85E-05 | 55.799

**Supplementary Fig. 12: Extended analysis of the endogenous neural activity recording from fig. 6.** The neural probe (fig. 2b) was mounted onto the microdrive and implanted into the mPFC of C57BL/6 wild type mice.

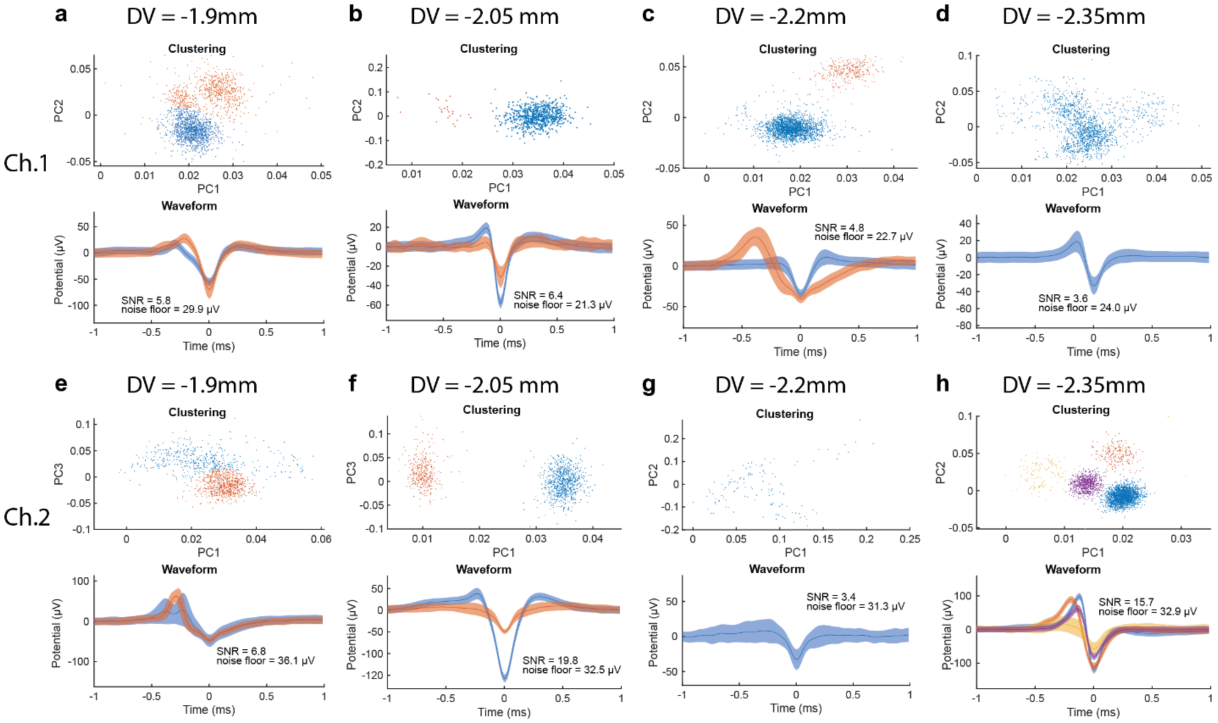


**Supplementary Fig. 13:** Analysis of the endogenous neural activity recording from the same mouse of fig.6, while the animal was awake. Similar to fig. 6, The implant was placed slightly above the mPFC at AP= +1.7 mm, ML= 0.4 mm, DV= -1.9 mm during the first day, and was lowered by 150 µm (1 turn) every day for 3 days. Endogenous neural activity was successfully recorded at each depth for channel 1 (e-h) and 2 (i-l) and putative single unit activity were isolated by principal-component analysis (top), and the average trace illustrates the waveform of the separated units (bottom).

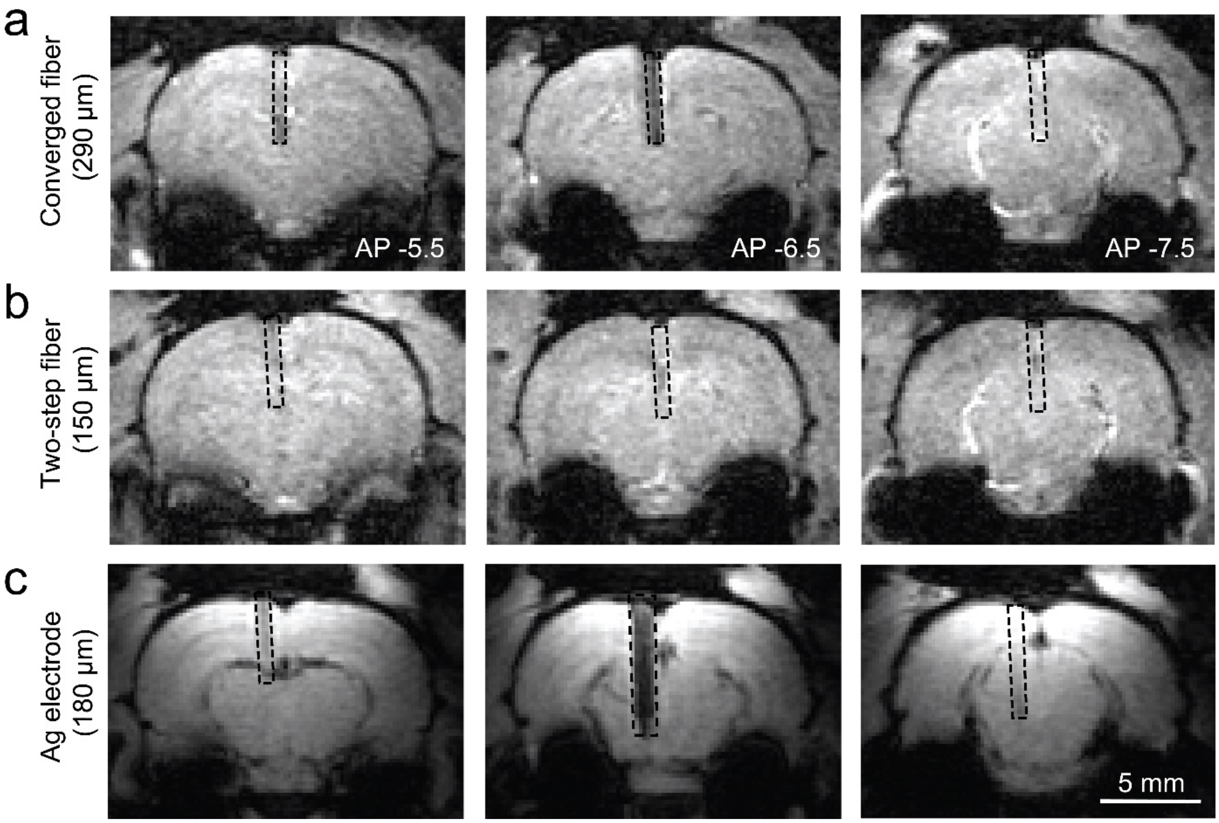


**Supplementary Fig. 14:** MRI image distortions caused by implants in comparison to silver wire which is routinely used for brain stimulation. Three consecutive slices of a T1-weighted fast low angle shot sequence with 1mm distance from AP -5.5 to AP -7.5 for convergence fiber (A) and two-step fiber (B), compared to a silver wire electrode (C) implanted in the rat brain.

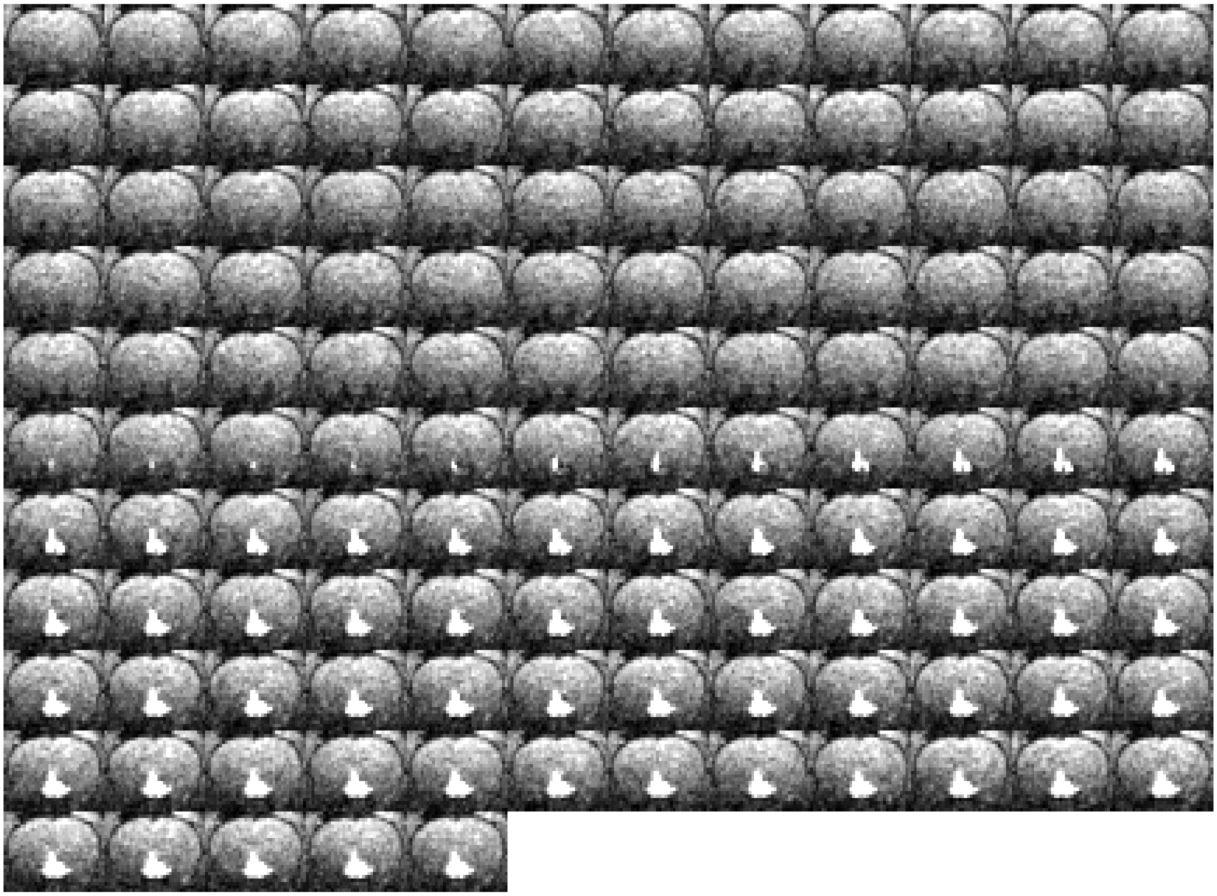

**Supplementary  Figure 15: A portfolio of snapshots acquired after every 10s during a typical infusion session with a two-step fiber.**

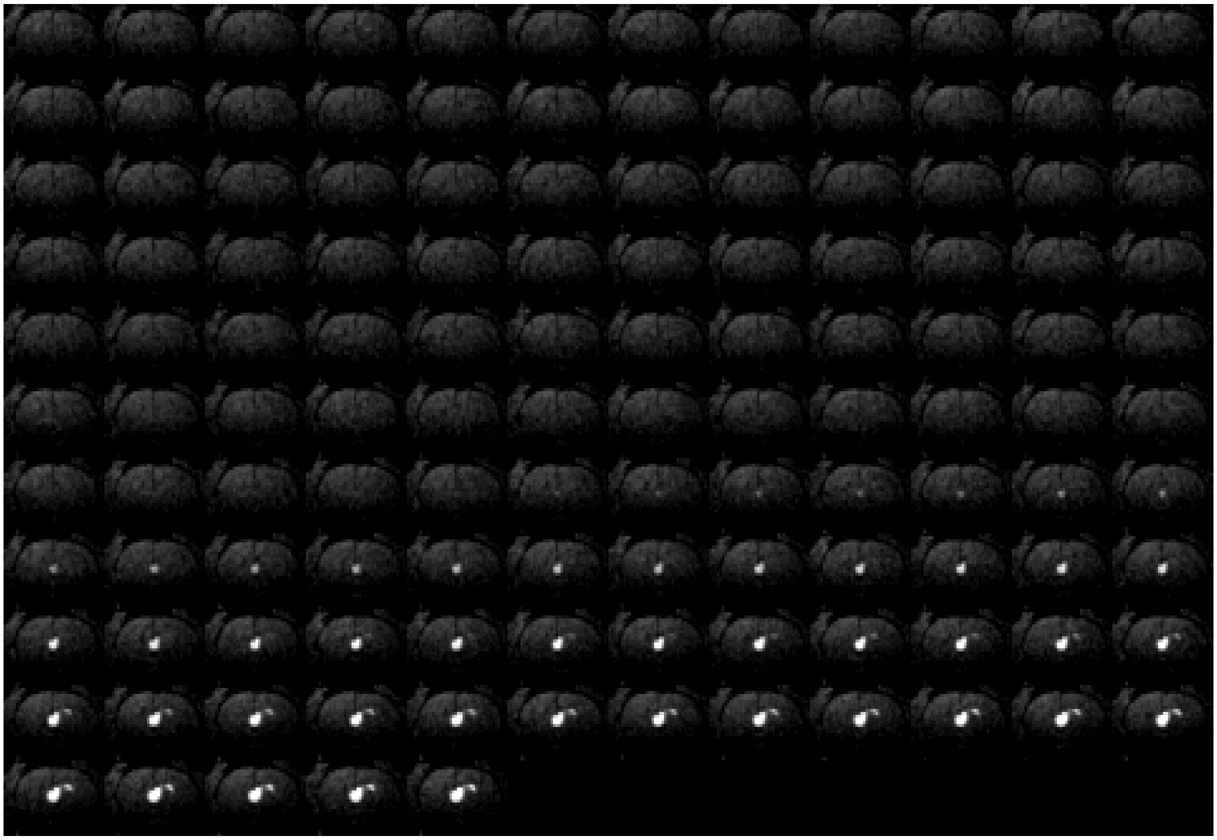

**Supplementary Figure 16:** A portfolio of snapshots acquired after every 10s during a typical infusion session with a converged multifunctional fiber.

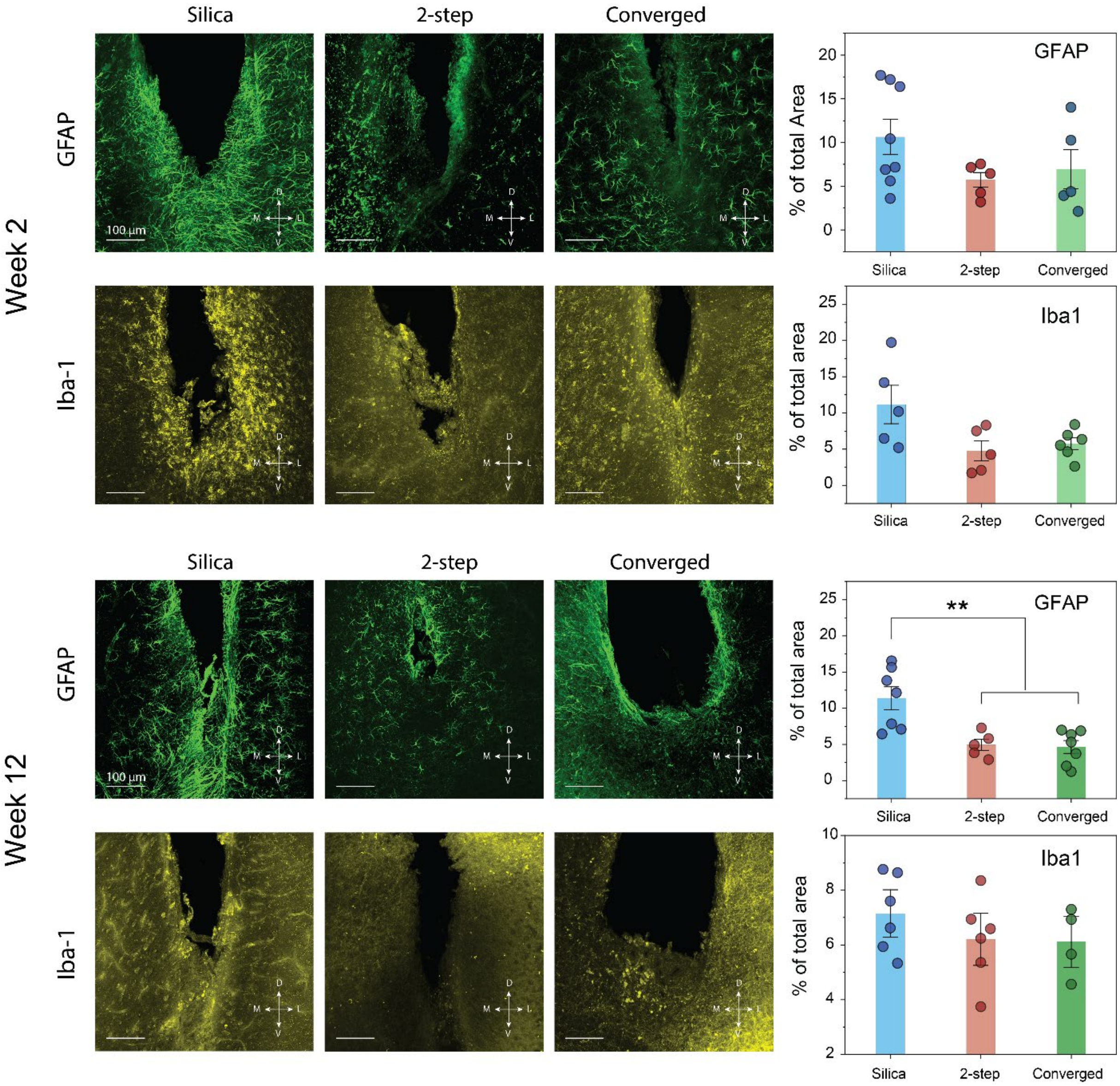


**Supplementary Figure 17:** Immunohistochemical evaluation of the foreign body response at 2- and 12-weeks post implantation. Scale bar = 100 µm.

**Supplementary Video 1:** Video of the convergence drawing.

**Supplementary Video 2:** Video showing real-time injection of a gadolinium-based MRI contrast agent Prohance (10mM) through an implanted two-step fiber.

**Supplementary Video 3:** Video showing real-time injection of a gadolinium-based MRI contrast agent Prohance (10mM) through an implanted converged-fiber.
